# Multiple cancer types rapidly escape from multiple MAPK inhibitors to generate mutagenesis-prone subpopulations

**DOI:** 10.1101/2023.03.17.533211

**Authors:** Timothy E. Hoffman, Chen Yang, Varuna Nangia, C. Ryland Ill, Sabrina L. Spencer

## Abstract

Many cancers harbor pro-proliferative mutations of the mitogen-activated protein kinase (MAPK) pathway and many targeted inhibitors now exist for clinical use, but drug resistance remains a major issue. We recently showed that BRAF-driven melanoma cells treated with BRAF inhibitors can non-genetically adapt to drug within 3-4 days to escape quiescence and resume slow proliferation. Here we show that this phenomenon is not unique to melanomas treated with BRAF inhibitors but rather is widespread across many clinical MAPK inhibitors and cancer types driven by EGFR, KRAS, and BRAF mutations. In all treatment contexts examined, a subset of cells can escape drug-induced quiescence within four days to resume proliferation. These escapee cells broadly experience aberrant DNA replication, accumulate DNA lesions, spend longer in G2-M cell cycle phases, and mount an ATR-dependent stress response. We further identify the Fanconi anemia (FA) DNA repair pathway as critical for successful mitotic completion in escapees. Long-term cultures, patient samples, and clinical data demonstrate a broad dependency on ATR- and FA-mediated stress tolerance. Together, these results highlight the pervasiveness with which MAPK-mutant cancers are able to rapidly escape drug and the importance of suppressing early stress tolerance pathways to potentially achieve more durable clinical responses to targeted MAPK pathway inhibitors.

## INTRODUCTION

The mitogen-activated protein kinase (MAPK) pathway is a major growth factor signaling force driving cellular proliferation. The protein cascade RAS-RAF-MEK-ERK facilitates this signaling, resulting in the activation of transcription factors that produce D-type cyclins and other cell-cycle effectors (*1*) (Fig. 1A). Cancer cells often harbor mutations in EGFR, KRAS, or BRAF that cause constitutive activation of the MAPK pathway and unfettered cellular proliferation. EGFR mutations are found in over 30% of all lung cancers (*2*), KRAS mutations in over 30% of lung adenocarcinomas (*3*) and up to 50% of colorectal cancers (*3*), and BRAF mutations in 5-12% of colorectal cancers (*4,5*) and over 50% of melanomas (*5*).

**Fig. 1.**
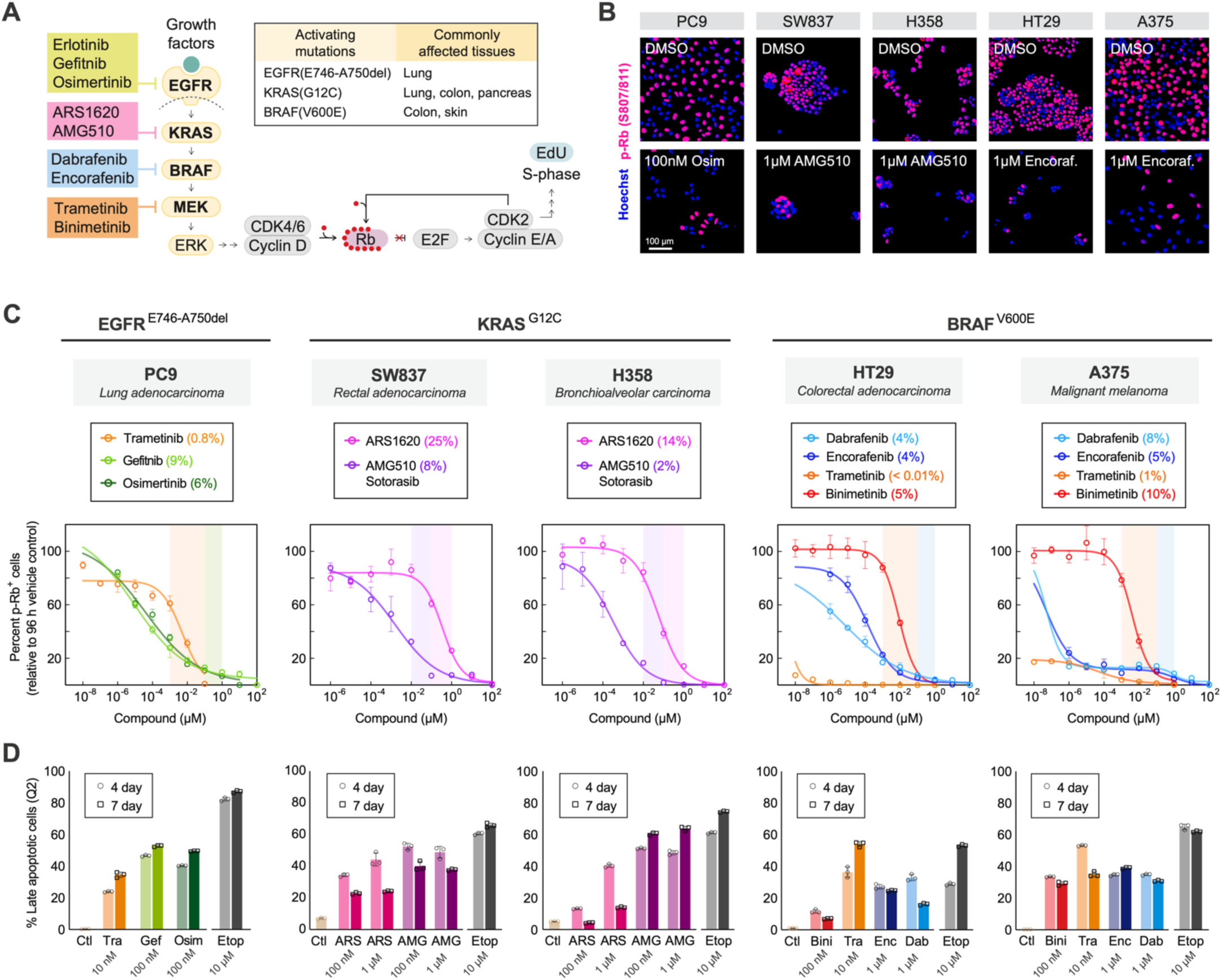
EGFR-, KRAS-, and BRAF-mutant cancer cells can survive and cycle through high-dose MAPK pathway inhibitors. **(A)** MAPK signaling pathway and subsequent cell-cycle progression. MAPK pathway inhibitors tested throughout this study are shown. **(B)** Representative immunofluorescence images of five human MAPK-mutant cancer cell lines 96 h after indicated treatments, stained for DNA content and phospho-Rb. **(C)** Dose-response profiling of MAPK-mutant cancer cell lines to their corresponding effective inhibitors after 96 h treatments. Fold change in phospho-Rb^+^ cells is normalized relative to 96 h DMSO condition. Shaded bars cover dose ranges commonly used in cell culture models. Reported % value in plot legends corresponds to the measured value remaining at the highest concentration of shaded bar regions. Mean ± std of 3 replicate wells. **(D)** Apoptosis measurements of five cancer cell lines in response to effective inhibitor doses, typically > IC_80_ from panel (C), reported as % of cells positive for propidium iodide and annexin by flow cytometry (see fig. S2, A and B, for gating scheme). Mean ± std of 3 replicates.

Small molecule cancer therapeutics that selectively target MAPK proteins have gained increasing clinical approval over the past two decades (*6*). Among the most successful drugs are EGFR, BRAF, and MEK1/2 inhibitors that offer high potency and selectivity for mutated forms of the proteins. First-generation ATP-competitive EGFR inhibitors erlotinib and gefitinib have similar clinical selectivity against mutant EGFR, while the newest inhibitor osimertinib offers the greatest potency yet for simultaneously inhibiting the initial EGFR driver mutations L858R and the common exon 19 deletion E746-A750del as well as the T790M mutation that is frequently acquired during erlotinib and gefitinib treatment (*7*). MEK non-ATP-competitive allosteric inhibitors trametinib, cobimetinib, and binimetinib offer anti-tumour efficacy in a variety of MAPK-mutant backgrounds (*8*). ATP-competitive BRAF inhibitors vemurafenib, dabrafenib, and encorafenib have also gained widespread approval for cancers harboring the common BRAF^V600E^ mutation, with encorafenib leading in bioavailability (*9,10*). KRAS inhibitors are the newest in development, as the RAS GTPase has been far more difficult to target than ATP-activated kinases (*6*). The potent AMG510 small molecule is the first clinically approved KRAS^G12C^-specific inhibitor (*11*), with another molecule ARS1620 and its derivatives still under investigation (*12,13*).

Despite the formidable progress of these compounds in the clinic, all are still subject to the threat of eventual resistance and tumour relapse (*14*). While drug resistance can be explained by expansion of rare subpopulations harboring pre-existing mutations or primed expression features (*15–18*), many reports point to the contributions of non-genetic drug adaptation mechanisms driving acquired resistance (*19–25*). This phenomenon has been identified in pivotal studies of acquired EGFR inhibitor resistance, where transient, non-genetic drug-induced states give rise to mutational evolution and diverse genetic resistance modes during extended treatment (*e.g.*, acquisition of T790M) (*19–21*).

In an effort to uncover the basis for rapid non-genetic drug adaptation, we recently discovered that subpopulations of melanoma cells escape drug-induced quiescence within the first 3 days of BRAF inhibitor treatment and continue to cycle slowly over weeks of treatment (*24*). We further found that this melanoma “escapee” subpopulation incurs DNA damage while cycling in drug. This is consistent with reports showing that subsets of BRAF-mutant colorectal cancer cells gain DNA damage within the first few days of dabrafenib and cetuximab treatment (*26, 27*). This DNA damage, in combination with low-fidelity DNA repair pathways, enables increased mutagenesis of cancer cells that survive in targeted therapy (*27*). Additionally, drug-induced DNA replication stress and chromosomal instability may also exacerbate the development of aneuploidy in escapee cancer cells, which can contribute to evolution of drug resistance (*28*). These reports highlight the need to identify ways to suppress the early drug adaptation that enables cells to escape drug, and also beg the question: how widespread is the phenomenon of rapid non-genetic escape from targeted therapies?

Here we tested whether rapid non-genetic escape from MAPK pathway inhibitors occurs in other cancer types beyond melanoma. We used multiple cancer types (colorectal adenocarcinoma, lung adenocarcinoma, melanoma) harboring mutations in either EGFR, KRAS, or BRAF that we treated with their corresponding targeted therapies. Using a variety of fixed and live-cell imaging approaches, we found that escapees indeed exist in all these contexts, and furthermore, that all these escapees broadly incur DNA lesions when cycling in the presence of these drugs. We find that this persistently cycling subpopulation experiences aberrant DNA replication, spends significantly longer time in G2-M cell cycle phases, and mounts an ATR-dependent stress response. Additionally, we identified that the downstream Fanconi anemia (FA) DNA repair pathway enables successful mitotic completion of escapees and prevents apoptosis. We find that DNA stress features are present in short-term (3-7 days) and long-term (2 months) MAPK inhibition treatment settings in culture, and are also present in patient tumour biopsies after treatment relapse. Thus, targeting early stress tolerance pathways may present an underappreciated strategy to eliminate adaptive escape from targeted therapy across multiple cancer types.

## RESULTS

### Multiple MAPK-mutant cancer types can escape high-dose MAPK pathway inhibitors

To assess the extent of the escapee phenomenon across different MAPK-mutants and tissue origins, we profiled five cell lines (PC9, SW837, H358, HT29, A375) with common activating mutations EGFR^E746-A750del^, KRAS^G12C^, and BRAF^V600E^ after treatment with a variety of targeted inhibitors (Fig. 1A and fig. S1A). Upon immunostaining for hyperphosphorylated Rb, a marker of cell-cycle commitment (*29*), we find that all cell lines contain a proliferative escapee subpopulation after 96 h of treatment (Fig. 1B). Considering that this may be an effect of sub-saturating MAPK inhibitor doses, we profiled all cell lines with wide dose-response ranges of multiple targeted therapies and found incomplete suppression of cycling cells even at supraphysiological drug concentrations (Fig. 1C and fig. S1, B to D). Among the commonly tested in vitro dose ranges that are considered clinically relevant, growth inhibition was measured at 60-90%, with no cases of 100% inhibition (fig. S1B). We chose to proceed with doses for each of the drugs that were largely effective in reducing the phospho-Rb population, representing concentrations that typically exceed the IC_80_ value. In the five MAPK-mutant cell lines tested, apoptosis levels were modest after 4 or 7 days of treatment across the inhibitor spectrum, with only 25-65% of cells presenting as apoptotic (Fig. 1D and fig. S2, A and B).

### Escapee cell cycles display DNA replication deficits and mitotic struggles

After identifying escape from drug-induced quiescence in multiple MAPK mutant cell types, we hypothesized that escaping cells may have irregular cell cycles due to the difficulty of cycling in drug. To begin, we surveyed DNA synthesis rates by way of EdU incorporation in cells treated with MAPK inhibitors for 96h and found that escapee cells never reached the same DNA replication rate as control cells (Fig. 2A). This reduced DNA synthesis rate was observed across PC9, SW837, H358, HT29, and A375 cell lines treated with multiple inhibitors (Fig. 2B), suggesting that cells escaping MAPK pathway inhibition broadly experience DNA replication stress (*30*). Hypothesizing that the rate of cell-cycle progression may be altered in escapees, we quantified all major cell-cycle phases in multiple cell lines following 4 days of targeted MAPK pathway inhibition. We found that cells are more likely to be found in G2 and M phases when cycling in the presence of drug (Fig. 2C and fig. S3, A to C), indicating prolonged time spent in later cell cycle phases, perhaps to recover from DNA replication insufficiencies (*31*).

**Fig. 2.**
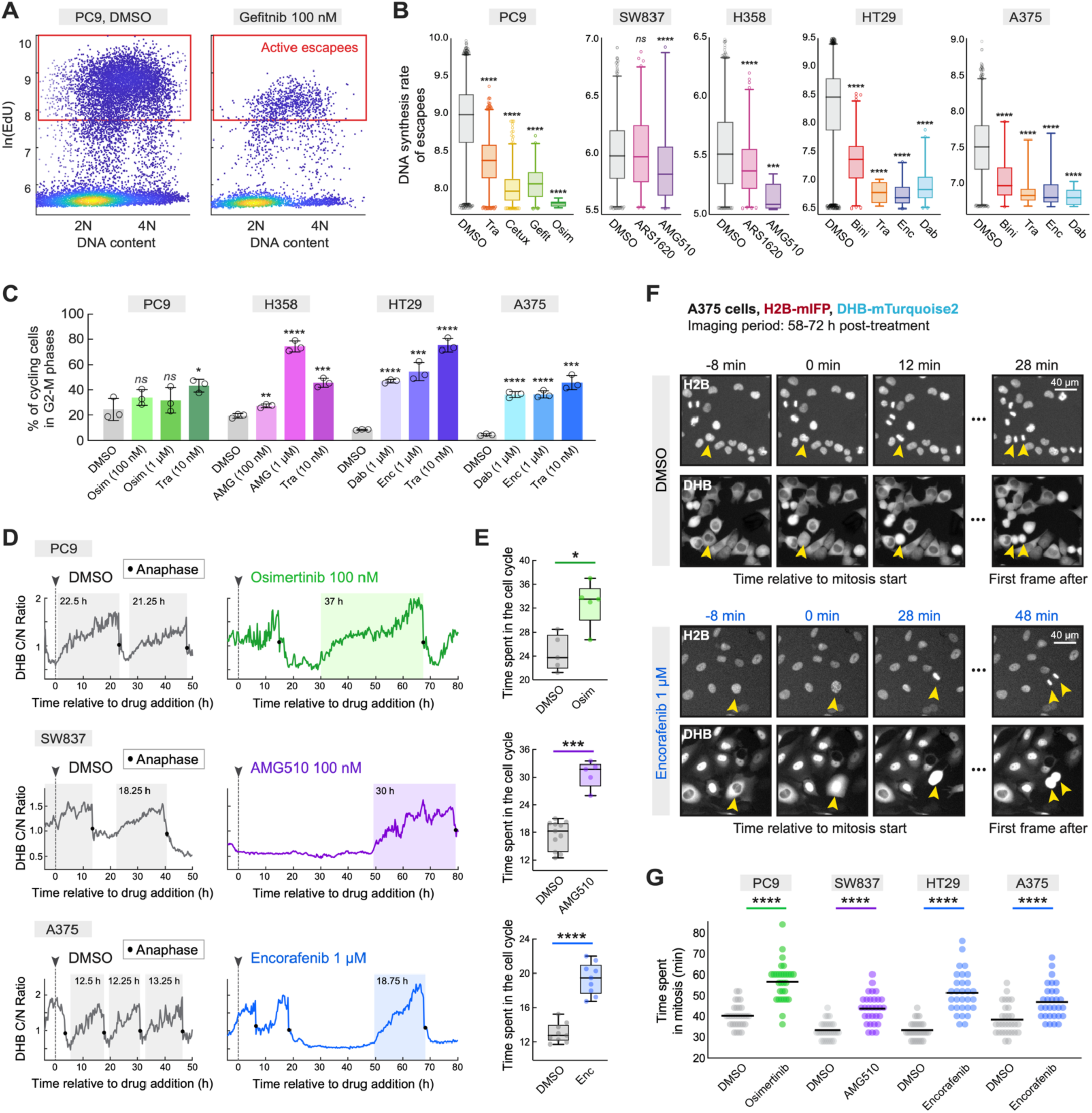
Escapee cell cycles display slowed DNA synthesis and increased time spent in G2-M. **(A)** Density scatter of DNA content and EdU incorporation in PC9 cells after indicated 96 h treatments. Red gate marks standard intensity of EdU+ control (n > 2000 cells). **(B)** Box plots of DNA synthesis rate in all cancer cell lines 96 h after indicated treatments. Box displays median and quartiles, whiskers display 1-99 percentile; *p*-values determined by Mann-Whitney U test. **(C)** Percent of cycling cells in G2-M after 96 h treatments (see fig. S3, A to C for gating scheme). Mean ± std of 3 replicates; *p*-value determined by unpaired t-test. **(D)** PC9, SW837, and A375 single-cell traces of DHB-based CDK2 activity sensor (see fig. S3D) over days in treatment. Time spent in active cell-cycle progression is highlighted by shaded bar based on time between CDK2 rise to anaphase. **(E)** Time spent in active cell-cycle progression (time from CDK2-rise to anaphase) for untreated proliferating cells and escapees. Box displays median and quartiles; *p*-values determined by Mann-Whitney U test. **(F)** Film strip of H2B and CDK2 activity sensor in A375 cells completing mitosis under DMSO treatment or BRAF inhibition. **(G)** Quantification of time spent in mitosis (from nuclear envelope breakdown to the point of anaphase) for indicated cell lines and MAPK inhibition treatments. n = 30 cells per condition; *p*-values determined by Mann-Whitney U test.

To study the cell-cycle dynamics in escapees, we tracked single-cell proliferation in real time over 4 days by time-lapse microscopy of a fluorescent biosensor for Cyclin-Dependent Kinase 2 (CDK2) activity (*32*) (fig. S3D). In PC9 EGFR-mutant lung cancer cells, SW837 KRAS-mutant colon cancer cells, and A375 BRAF-mutant melanoma cells treated with high doses of osimertinib, AMG510, and encorafenib, respectively, the drug initially caused cells to enter a CDK2-low quiescence as expected. However, after variable amounts of time in this state, a subpopulation of escapees emerged, marked by a return to the CDK2-increasing trajectory (Fig. 2D, fig. S3E and Video S1). Upon further inspection of the full escapee cell cycle by this method, we noted that cell-cycle durations of all MAPK-mutant escapees were significantly lengthened relative to those of vehicle-treated cells (Fig. 2, D and E), consistent with the idea that escapees are struggling through the cell cycle in the presence of drug. By performing time-lapse experiments with a faster frame rate, we also observed longer mitotic intervals in escapees compared with control cells (Fig. 2, F and G, fig. S4, A to C, Video S2 and Video S3), a potential consequence of DNA under-replication causing chromatin segregation problems (*33, 34*). These observations highlight the need to understand how cancer cells escaping potent MAPK pathway inhibition can survive and cycle slowly in the face of these cell-cycle irregularities.

### MAPK-mutant escapees incur DNA damage and mount DNA replication stress responses

As escapee cells appear to labor through the cell cycle in drug, we tested whether these cells show signs DNA damage (Fig. 3A). In comparison to unperturbed cycling cells, PC9, SW837, H358, HT29 and A375 escapees cycling in the presence of MAPK inhibitors displayed significantly more intense phospho-H2AX S139 (γ-H2AX) staining, a marker of DNA breaks, relative to non-escapees (*35*) (Fig. 3, B to D). As another readout of DNA damage, we measured expression of the p21 CDK inhibitor, which is known to be upregulated by p53 in situations of DNA replication stress and prolonged mitosis (*36,37*). A375 cells (which are p53-competent) showed upregulation of p21 over the course of 2 days of BRAF inhibition (fig. S5A).

**Fig. 3.**
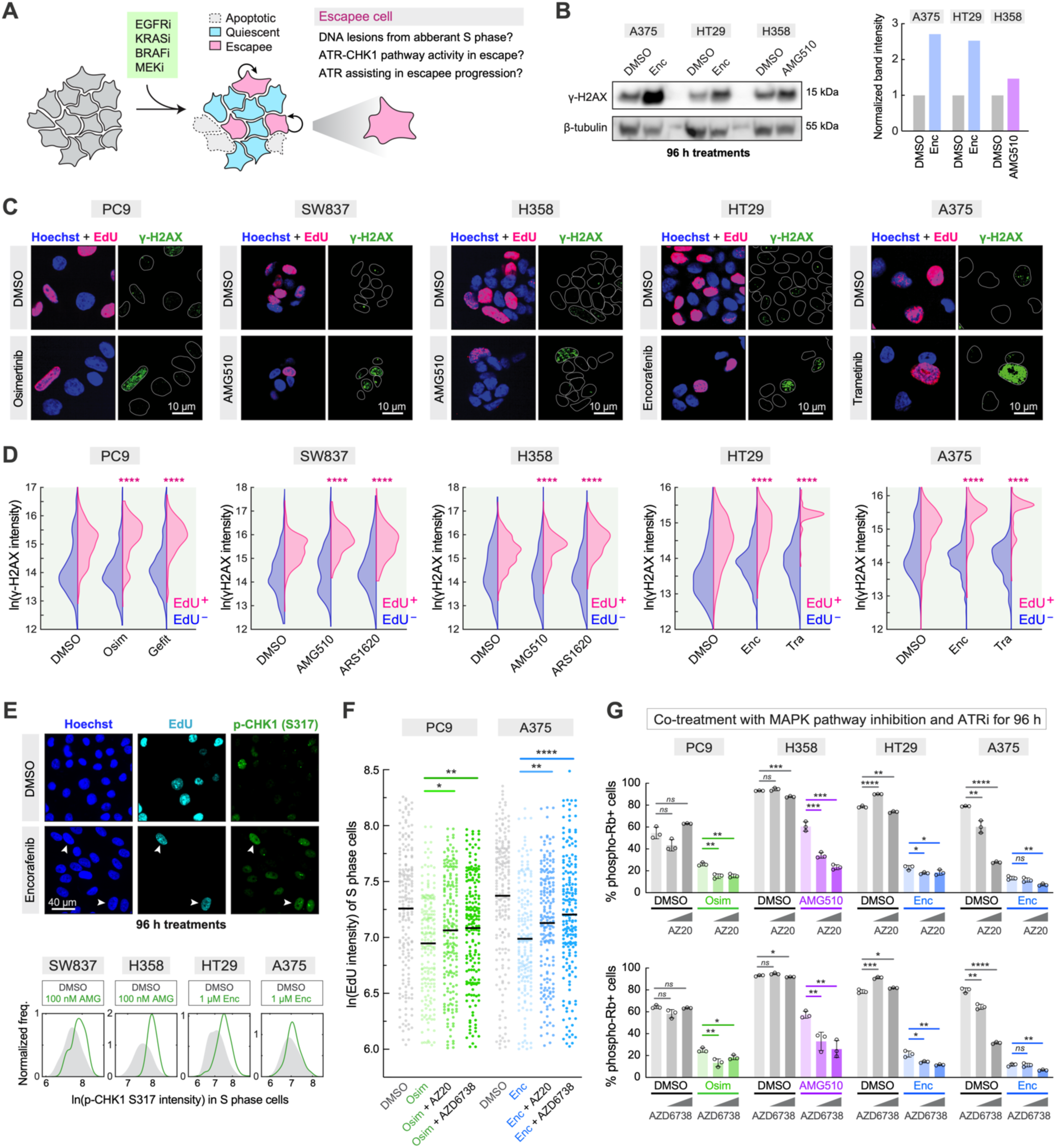
Cells escaping MAPK inhibition show widespread indications of DNA damage accumulation and DNA replication stress responses. **(A)** Illustration of the hypothesis that escapees mount a replication stress response to manage aberrant S phases and tolerate DNA lesions. **(B)** Western blot analysis of γ-H2AX in the indicated cell lines after 96 h treatments. β-tubulin is used as a loading control. Bar plot displays quantification of γ-H2AX band intensity normalized to respective loading control band, and values are reported as fold change of control. **(C)** Representative immunofluorescence images of five human MAPK-mutant cancer cell lines after 72 h indicated treatments, stained for DNA content, EdU incorporation, and γ-H2AX. **(D)** Quantification of γ-H2AX nuclear fluorescence intensity after 72 h indicated treatments in EdU^−^ and EdU^+^ cells, displayed as split violins; *p*-values determined by Mann-Whitney U tests of MAPK-inhibited EdU^+^ cells compared with DMSO-treated EdU^+^ cells. (E) *Top:* Representative immunofluorescence images of phospho-CHK1 (S317) in A375 cells escaping BRAF inhibition (cycling status displayed by EdU intensity). Bottom: Histograms of phospho-CHK1 (S317) nuclear intensity in S phase cells (EdU^+^) in the indicated cell lines after 96 h MAPK inhibition. **(F)** Distribution of EdU intensity in escapees (after 100 nM Osimertinib or 1 μM Encorafenib) co-treated with ATR inhibition (100 nM AZ20 or 100 nM AZD6738). 96 h treatments. Line drawn at the mean value; n = 190 cells plotted per condition; *p*-values determined by Mann-Whitney U test. **(G)** Quantification of cycling cells (determined by phospho-Rb S807/811 immunofluorescence) after 96 h of indicated treatment combinations with ATR inhibitors. AZ20 doses: 100 nM, 1 μM. AZD6738 doses: 100 nM, 1 μM. Mean ± std of 3 replicates; *p*-value determined by unpaired t-test.

Since γ-H2AX is a substrate of the ataxia telangiectasia and RAD3-related (ATR) master replication stress kinase (*38*), and since ATR can suppress DNA replication rates (*30*), we tested whether this pathway is activated in the escapee subpopulation (Fig. 3A). To address this, we first immunostained for phosphorylated ATR and phosphorylated checkpoint kinase 1 (CHK1), a canonical downstream target of ATR (*39*), and found that these markers were upregulated in escapees in all cell lines and drugs tested (Fig. 3E and fig. S5, B and C). To test whether the ATR-CHK1 pathway suppresses DNA replication rates in escapees, we co-treated PC9 and A375 cells with their relevant MAPK inhibitors combined with selective ATR inhibitors AZ20 or AZ6738 and found that DNA replication rates can indeed be partially rescued with this combination (Fig. 3F). Next, we co-treated PC9, H358, HT29 and A375 cells with their relevant MAPK inhibitor and ATR or CHK1 inhibitors for 96 h and found fewer escapees, as indicated by up to 2.5X fewer phospho-Rb positive cells (Fig. 3G and fig. S5D). These results indicate that pervasive replication stress across escapees must be responded to by an ATR-dependent mechanism in order to successfully progress through a drugged cell cycle.

We also tested whether the ataxia telangiectasia mutated (ATM) kinase is also active in the escapee subpopulation, since this pathway recognizes double-strand breaks and shares some cooperativity with ATR activity (*38*). Indeed, co-treating PC9, H358, HT29 and A375 cells with MAPK pathway inhibition and clinical ATM inhibitor AZD0156 resulted in fewer escapees relative to MAPK inhibition alone (fig. S5E). Finally, we measured apoptotic cell death after co-targeting the MAPK pathway and ATR for 7 days and found that this combination therapy results in cooperative killing of PC9, H358 and A375 cells (fig. S5, F and G). Together, these data indicate that MAPK-mutant cells escaping targeted therapy mount ATR- and ATM-directed stress response programs that broadly help them tolerate the DNA replication stress and DNA damage they experience while cycling in drug, and that inhibiting these pathways reduces the fraction of early escapees via apoptosis.

### MAPK-mutant escapees broadly depend on FA-mediated DNA replication recovery

The FA complementation group constitutes a conserved pathway for repairing DNA interstrand crosslinks and for general protection of DNA replication fork function under stress (*40–45*). The pathway is commonly activated in response to ATR activity (*38*), which allows for formation and ubiquitination of the FANCD2/I heterodimer that clamps onto DNA lesions and recruits the downstream elements required for excision, bypass, and repair.

To test whether this pathway is involved in escapee’s tolerance of abnormal cell cycles, we first performed immunofluorescence imaging of PC9, H358, SW837, HT29 and A375 cells after 72 h of MAPK pathway inhibition and found that that EdU-positive escapees display increased nuclear FANCD2 (Fig. 4, A to C). By quantifying the relative FANCD2 intensity in all major cell cycle phases, we noted increased intensity in late S and G2/M phases (Fig. 4B). To precisely define the cell-cycle dynamics of FANCD2 recruitment, we created an A375 cell line expressing fluorescently tagged FANCD2-mCitrine (*46*), along with the DHB-mCherry CDK2 sensor and H2B-mIFP for cell tracking (fig. S6A). We filmed these cells for 72 h in the presence of dabrafenib and noted the striking appearance of intense nuclear foci in the final hours of the cell cycle in escapees (Fig. 4, D and E, fig. S6B and Video S4). This timing is consistent with a role for FANCD2 toward the end of DNA replication and through mitosis under unique stress conditions (*34,47*), suggesting that MAPK-mutant cancer cells escaping targeted therapies do not properly complete DNA replication and may enter mitosis with under-replicated regions of the genome.

**Fig. 4.**
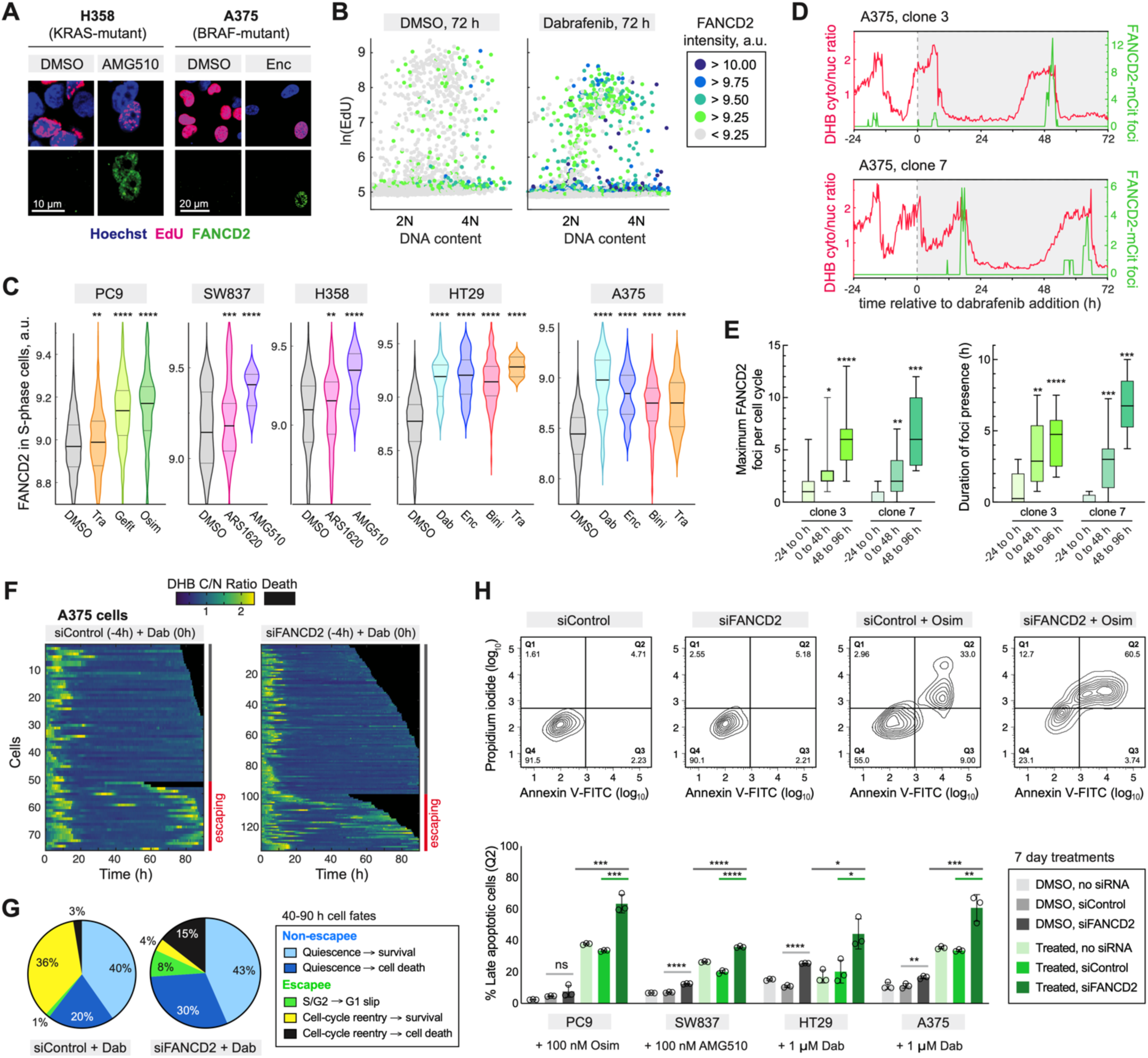
Escapee cancer cells broadly rely on the recruitment of FANCD2 in late S/G2 to successfully complete mitosis in drug. **(A)** Representative immunofluorescence images of cells after 72 h of indicated treatments, stained for DNA content, EdU incorporation, and FANCD2. **(B)** Scatter plots of DNA content and EdU intensity in A375 cells treated with 72 h DMSO or BRAF inhibition. Each data point is color-coded to represent nuclear FANCD2 intensity. **(C)** Quantification of FANCD2 nuclear fluorescence intensity after 72 h indicated treatments in EdU^+^ cells, displayed as violins; *p*-values determined by Mann-Whitney U tests of MAPK-inhibited cells compared with DMSO-treated cells. **(D)** Representative single-cell traces of A375 cells expressing both the CDK2 activity sensor and FANCD2-mCitrine tag. FANCD2-mCitrine dynamics are quantified as the number of nuclear foci that appear. **(E)** Quantification of maximum FANCD2 foci number or duration for untreated cells (−24 to 0h) or cell cycles spanning early or later treatment periods. n > 12 cells per condition; *p*-values determined by Mann-Whitney U test. **(F)** Heatmaps of single-cell CDK2 activity traces of A375 cells after indicated treatments and siRNAs. Each row represents the CDK2 activity in a single cell over time according to the colormap. Traces that end due to cell death are marked by black coloring. **(G)** Fate summary of cells observed over the 40-90 h imaging period from the experiment in panel (F). **(H)** Flow cytometric apoptosis assay in MAPK-mutant cell lines after the indicated 7-day treatments. *Top panels*, Representative contour plots of PC9 cells progressing toward apoptosis (quadrant Q2) after EGFR inhibition with 100 nM Osimertinib and FANCD2 depletion. *Bottom panel*, Quantification of Q2 across indicated cell lines and treatments. Mean ± std of 3 replicates; *p*-value determined by unpaired t-test.

We next tested whether FANCD2 is necessary for successful completion of mitosis under drug pressure. After depleting FANCD2 by siRNA in A375 cells (fig. S6C), we performed time-lapse imaging under BRAF inhibition. Relative to the siControl-treated cells, FANCD2-deficient A375 cells experienced 9-fold lower incidence of escape with mitotic completion and 2-fold increased total incidence of cell death (Fig. 4, F and G, fig. S6D and Video S5). It is important to note that both escapees and non-escapees experienced increased cell death after FANCD2 depletion, suggesting that the non-escapees may have gained new unresolvable damage in their final cell cycles before entering MAPKi-induced quiescence. In escapee cell cycles, FANCD2 deficiency resulted in many more instances of death during mitosis and cases of mitotic slippage (*i.e.*, movement from G2 to G0/G1 with no cell division) (Fig. 4G). We confirmed the importance of FANCD2 for successful cell-cycle completion in drug across other EGFR-, KRAS-, and BRAF-mutant cell lines, finding that concurrent FANCD2-deficiency and MAPK pathway inhibition resulted in cooperative killing of cancer cells (Fig. 4H). Some cell lines experienced modest yet significant increases in cell death after FANCD2 depletion alone (Fig. 4H), suggesting that MAPK-mutant cancer cells may be broadly dependent on this DNA recovery mechanism. These observations are consistent with other reports demonstrating the ability of the FA pathway to maintain proliferation and survival, including in BRAF-mutant melanomas (*48,49*).

To understand the broad stress tolerance in MAPK-mutant cancers escaping targeted MAPK inhibitors, we next investigated low-fidelity polymerase kappa (Polκ) induction, which is known to cooperate with the FA pathway to facilitate DNA replication fork restart after instances of stress-induced stalling (*50*). To measure Polκ induction, we quantified the translocation of this enzyme from the cytoplasm to the nucleus (*51*). By immunofluorescence, we observed a sustained translocation of Polκ to the nucleus in A375 cells after treatment with dabrafenib, encorafenib, binimetinib, or trametinib (fig. S7, A and B), consistent with reports of melanoma and lung cancer cells treated with vemurafenib and erlotinib, respectively (*51*). We also observed this drug-induced translocation in both escapees and non-escapees across PC9, SW837, H358, HT29 and A375 cells upon treatment with drugs that target their respective MAPK drivers (fig. S7, C and D). Taken together, these data underscore a unique stress tolerance of cells escaping MAPK pathway inhibitors carried out by the FA pathway and low-fidelity polymerase induction.

### Long-term cultures and clinical samples highlight broad replication stress tolerance features of escapees

To determine the importance of the ATR-mediated stress response upon extended MAPK pathway inhibition, we treated EGFR-mutant, KRAS-mutant and BRAF-mutant cells with their appropriate targeted MAPK inhibitors for 2 months, with or without co-treating with a clinical ATR inhibitor. Indeed, cells receiving this long-term combination therapy had significantly less growth compared to cells treated with MAPK inhibition alone (Fig. 5A). Similarly, cells co-treated with MAPK pathway inhibition and a clinical ATM inhibitor also had significantly less growth compared to cells treated with MAPK inhibition alone (fig. S8A).

**Fig. 5.**
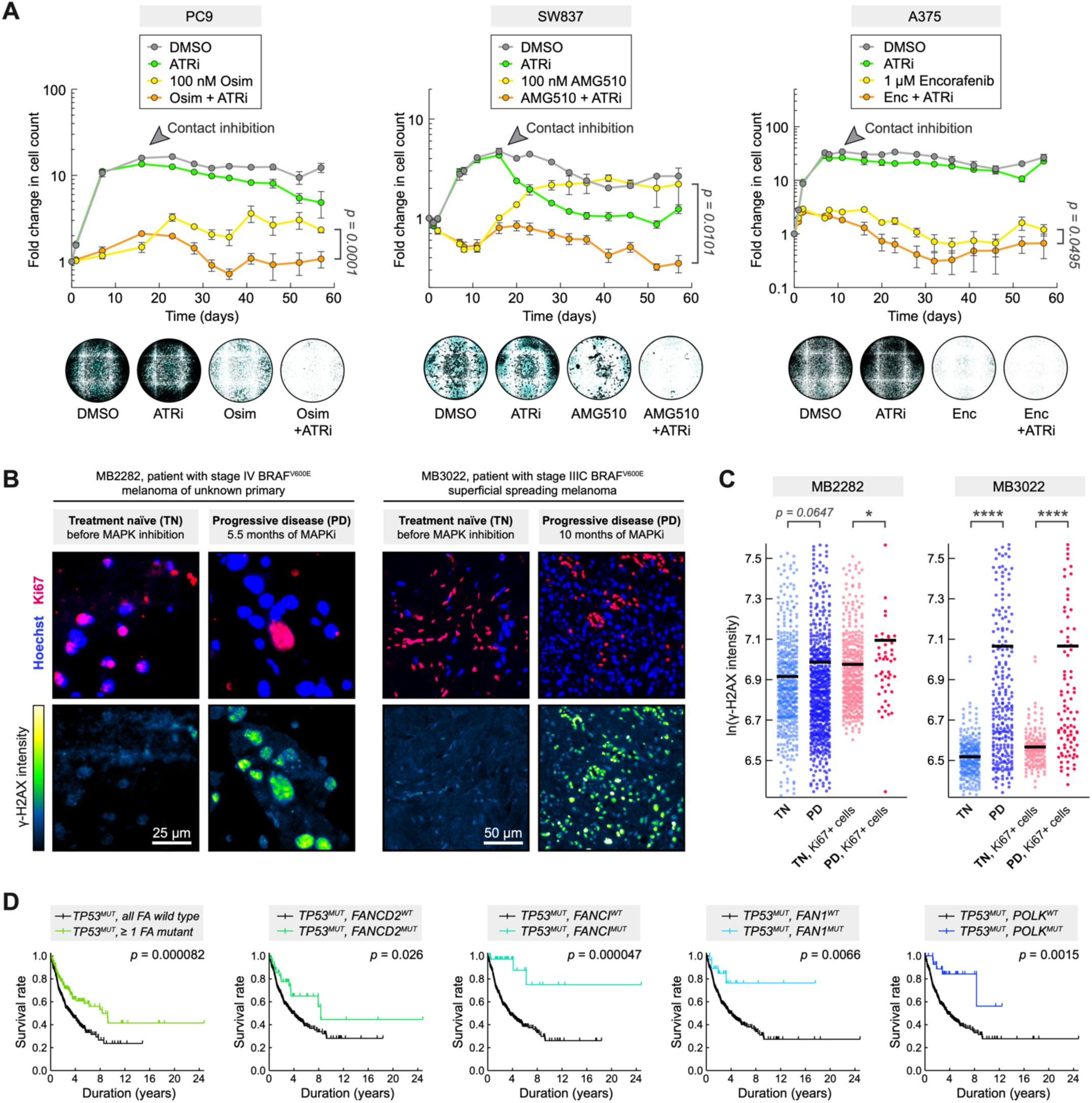
Long-term cultures, patient tumours and clinical data highlight replication stress tolerance features of escapees in extended treatment. **(A)** *Top:* Long-term culture of MAPK-mutant cell lines with indicated MAPK inhibitors and clinical ATR inhibitor (5 μM AZD6738). Cell count was quantified longitudinally by imaging H2B-mIFP fluorescence. Unfettered proliferation in control groups causes contact inhibition within first 2 weeks of imaging. Mean ± std of 4 replicate wells; *p*-value determined by unpaired t-test of final time point. *Bottom:* Representative wells from the final day of imaging. Inverted H2B-mIFP fluorescence shown. **(B)** Formalin-fixed paraffin-embedded (FFPE) tissue sections longitudinally biopsied before and after treatment from patients with BRAF^V600E^ melanoma (see fig. S8B for patient and tumour details). Immunofluorescence of Ki67 (proliferation marker) and γ-H2AX (DNA damage marker) displayed. **(C)** Quantification of all cells (shown in blue coloring) and Ki67-positive cells (shown in red coloring) in FFPE sections with elevated γ-H2AX intensity. Line drawn at the mean; n > 235 cells scored before Ki67 gating; p-value determined by Mann-Whitney U test. **(D)** Kaplan-Meier survival analyses based on data generated from the Harmonized Cancer Datasets within the Genomic Data Commons (GDC) research database (as of April 28, 2022; https://portal.gdc.cancer.gov). Patients were included with tumours harboring ≥ 1 EGFR, KRAS, or BRAF mutation and ≥ 1 TP53 mutation. Cohort comparisons were made between tumour subgroups containing wild-type or mutant forms of the indicated genes. Left-most plot examines a cohort that has ≥ 1 mutation among a list of FA-associated genes (see fig. S9A for FA gene list). Displayed *p*-values were determined by log-rank test.

To assess the escapee phenomenon in a clinical context, we obtained patient samples of BRAF^V600E^ melanoma tumours longitudinally biopsied before treatment and after cancer recurrence (fig. S8B). In these tissue sections, we co-stained for Ki67 to measure proliferative activity (*52*) and γ-H2AX. We found that Ki67-positive cells that exist after extended BRAFi and MEKi combination treatment (escapees) had significantly higher levels of γ-H2AX relative to Ki67-positive cells from samples prior to MAPK inhibition (Fig. 5, B and C). These observations indicate that patient cancer cells that cycle in drug also incur DNA damage in vivo, pointing to increased mutagenesis and increased potential for acquisition of drug-resistance mutations.

We also analyzed tumour mutation profiles relative to patient survival data from the Harmonized Cancer Datasets within the Genomic Data Commons (GDC) research database and found that cases with at least 1 targetable MAPK mutation in a *TP53*-mutant background had significantly higher survival rates if the tumours also harbored any variety of FA pathway or *POLK* mutations (Fig. 5D, fig. S9A). MAPK-mutant cases with intact *TP53* also experienced higher survival rates, though not always to a significant extent for the different FA pathway mutants (fig. S9B). These results are consistent with our findings that a defective FA pathway can prevent successful mitoses of tumour cells under drug pressure (Fig. 4, F and G), especially in common genetic backgrounds of p53 functional loss that cause unchecked tumour growth (*53*). Four out of five cell lines used in our study harbor *TP53* mutations (fig. S1A), and all show dependence on ATR and FA pathway activity to successfully escape targeted MAPK pathway inhibition.

## DISCUSSION

The phenomenon of rapid non-genetic escape from targeted therapy is coming to the forefront of cancer drug resistance research, and the work presented here highlights just how broad this phenomenon is across cancer cell types treated with targeted MAPK pathway inhibitors. Cancer cell “persisters” have been described as rare cells that merely survive treatment, whereas we refer to cells that survive *and* slowly cycle during treatment as “escapees” and find that they are not particularly rare. The cycling activity of the escapee subpopulation, combined with tolerable DNA damage accumulation, yields a reservoir of cells with high potential for new mutagenesis events and acquired genetic drug resistance (Fig. 6).

**Fig. 6.**
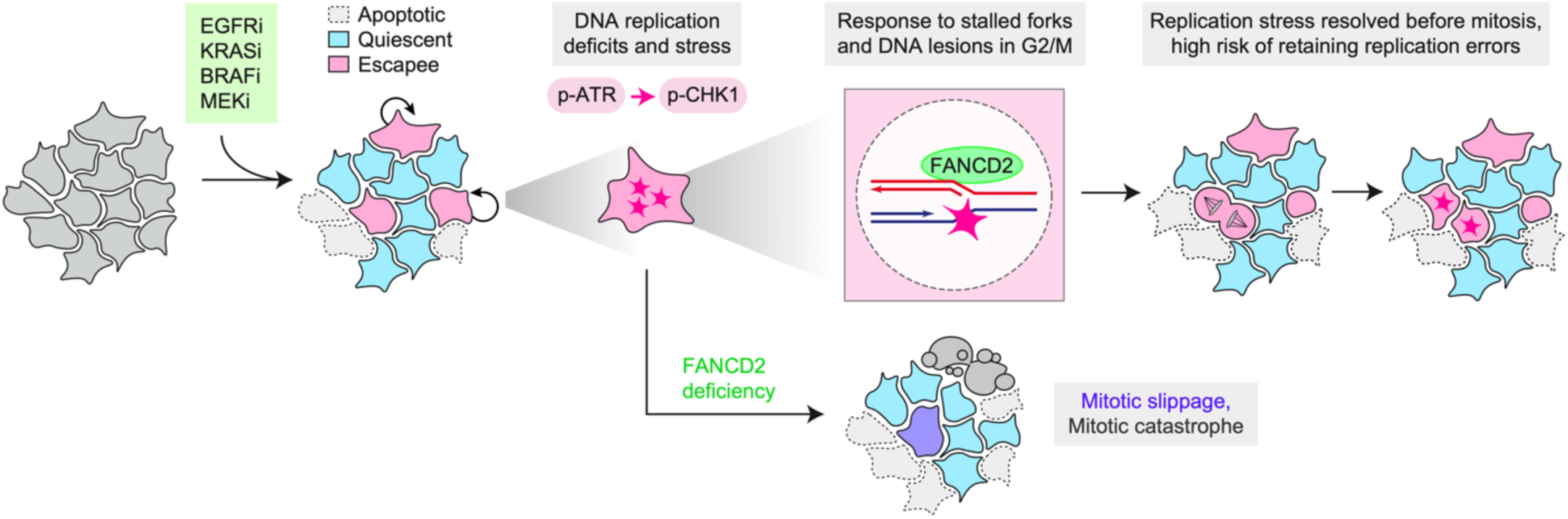
Proposed model of early drug escape, stress tolerance, and cancer cell evolution in extended treatment. Cells treated with drug have divergent fates within the first few days of treatment, where they will undergo cell death, remain quiescent, or escape treatment and re-enter the cell cycle. These escapee cells will suffer DNA replication stress, relying on the FA pathway to restore replication fork stability and complete mitosis.

Among the most curious effects of the MAPK pathway inhibitors tested here is their ability to induce DNA damage, since these drugs were not intended to have any direct genotoxicity. Intuitively, the increases in DNA damage must be indirect, likely stemming from cells being forced through cycles of DNA replication without sufficient preparation or resources. Because FANCD2 is known to respond to nucleotide depletion (*41*), it is possible that escapee cells have not prepared ample nucleotide pools during their periods of quiescence leading up to the point of cell-cycle re-entry, since these pools are partially dependent on MAPK signaling (*54,55*). Escapee cells also appear to rush through their G1 phase with insufficient time to license origins of replication (*24*), which could exacerbate genomic instability in the long term (*56*). Another possible source of DNA damage in escapees is oncogene-induced hyper-transcription, where hyperactive MAPK signaling can cause overwhelming transcriptional bursts and ATR activation (*57*). Identifying the root cause(s) of this DNA damage will help in developing strategies to push DNA damage in escapees out of the tolerable range and thereby push escapees toward apoptosis (*58*).

We and others have emphasized the FA pathway and related effectors (*e.g.*, ATR, Polκ) as an attractive set of targets for inducing synthetic lethality in tumours (*40, 59*). Here we show that targeting FA proteins themselves reduces the number of MAPK-mutant escapees in EGFR- KRAS- and BRAF-mutant cancers. Since the surviving escapee population can be reduced by 9-fold by co-inhibiting the FA pathway simultaneously with targeted MAPK inhibition, this type of polypharmacology approach may be effective as a transient or intermittent treatment to reduce survival of escapees. While the FA pathway has been valued as essential for somatic cells, low-dose, short or episodic treatments may greatly reduce the toxicity seen in long-term or permanent FA pathway ablation. Several small molecules have been developed that target FANCM, FANCL, and Polκ (*59*), though they are still in preclinical stages and have not yet been thoroughly investigated. If the FA pathway cannot be directly targeted even in short treatment intervals, targeting upstream kinases also offer promise. Clinical ATR and ATM inhibitors already exist for a variety of indications, and our work suggests that both of these master kinases are at play in helping many types of MAPK-mutant escapee cells adapt to cell-cycle stress and DNA damage, paralleling work noting the effective co-treatment of MAPK and ATM inhibitors (*60*).

In conclusion, our observations demonstrate the broad applicability of the escapee phenotype across MAPK-mutant cancers and targeted MAPK therapies, and suggests that suppressing the DNA replication stress tolerance of these early subpopulations may help achieve more durable clinical responses to targeted MAPK pathway inhibitors.

## MATERIALS AND METHODS

### Cell culture and cell line generation

The SW837 KRAS^G12C^ colorectal adenocarcinoma (#CCL-235), the H358 KRAS^G12C^ lung adenocarcinoma (#CRL-5807), the HT29 BRAF^V600E^ colorectal adenocarcinoma (#HTB-38) and the A375 BRAF^V600E^ malignant melanoma (#CRL-1619) cell lines were purchased from American Type Culture Collection (ATCC). PC9 cells were purchased from MilliporeSigma (#90071810). A375 cells were cultured at 37 °C with 5% CO2 in DMEM (Thermo Fisher, #12800-082) supplemented with 10% FBS, 1.5-g/L sodium bicarbonate (Fisher Chemical, #S233-500) and 1X penicillin/streptomycin. PC9, SW837, H358, and HT29 cells were maintained at 37 °C with 5% CO2 in RPMI1640 (Thermo Fisher, #22400-089) supplemented with 10% FBS, 1X Glutamax, 1X sodium pyruvate (Thermo Fisher, #11360-070) and 1X penicillin/streptomycin.

Low-passage PC9, SW837, HT29 and A375 cells were transduced with H2B-mIFP and DHB-mTurquoise2 lentivirus, that was prepared as described previously (*32*). Cells stably expressing these sensors were isolated by two rounds of FACS. A375 cells were transduced with FANCD2-mCitrine lentivirus, prepared with the pLenti-WT-FANCD2-mCitrine plasmid (*46*) that was a gift from Dr. Jean-Yves Masson at Laval University Cancer Research Center. A375 FANCD2-mCitrine single clones were isolated by FACS, validated by localization of nuclear foci after hydroxyurea treatment, and stably transduced thereafter with H2B-mIFP and DHB-mCherry lentivirus to create the final 3-color cell line.

### Small molecules

Treatments used in this study include: ZD1839 (gefitinib, EGFRi, Selleckchem #S1025); AZD9291 (osimertinib, EGFRi, Selleckchem #S7297); C225 (cetuximab, EGFRi, Selleckchem #A2000); AMG510, (sotorasib, RASi, Selleckchem #S8830); ARS1620 (preclinical RASi, Selleckchem #S8707); GSK2118436 (dabrafenib, RAFi, Selleckchem #S2807); LGX818 (encorafenib, RAFi, Selleckchem #S7108); GSK1120212 (trametinib, MEKi, Selleckchem #2673); MEK162 (binimetinib, MEKi, Selleckchem #S7007); VP16 (etoposide, TOP2i, Selleckchem #S1225); AZ20 (ATRi, Selleckchem #S7050); AZD6738 (ceralasertib, ATRi, Selleckchem #S7693); LY2603618 (rabusertib, CHK1i, Selleckchem #2626); AZD0156 (ATMi, MedChemExpress #HY-100016); Aphidicolin (DNA Pol α/δ inhibitor, Cell Signaling Technology #32774), NSC32065 (hydroxyurea, RNRi, Selleckchem #1896); PD0332991 (palbociclib, CDK4/6i, Selleckchem #S1116); AY22989 (rapamycin, mTORi, Selleckchem #S1039). Doses of MAPK inhibitors used throughout the study, unless otherwise specified, are: osimertinib, 100 nM; gefitinib, 100 nM; cetuximab, 100 μg/mL; ARS1620, 100 nM; AMG510, 100 nM; encorafenib, 1 μM; dabrafenib, 1 μM; trametinib, 10 nM; binimetinib, 100 nM.

### siRNA transfection

siRNA transfections were performed using the DharmaFECT 4 reagent (Dharmacon, #T-2004-02) according to the manufacturer’s instructions. The transfection mix was added to the cells 4 h prior to the time of drug treatment and removed 4 h after the time of drug treatment. Knockdown efficiency was determined by western blotting 24 h post-transfection. Oligonucleotides used in this study are: DS NC-1 (IDT, #51-01-14-04) and *FANCD2* DsiRNA (IDT, #hs.Ri.FANCD2.13.1, #hs.Ri.FANCD2.13.2, #hs.Ri.FANCD2.13.3; all were equally pooled for FANCD2 knockdown).

### Antibodies

Primary antibodies used in this study include: phospho-Rb (S807/811) (D20B12) rabbit mAb (1:500, Cell Signaling Technology #8516); phospho-Rb (S780) mouse mAb (1:1000, BD Biosciences #558385); phospho-Histone H3 (S10) (D2C8) rabbit mAb (1:400, Cell Signaling Technology #3377); phospho-Histone H3 (S10) (6G3) mouse mAb (1:400, Cell Signaling Technology #9706); phospho-H2AX (S139) (20E3) rabbit mAb (1:400 for standard immunofluorescence, 1:100 for FFPE tissue staining, Cell Signaling Technology #9718); p21 (12D1) rabbit mAb (1:400, Cell Signaling Technology #2947); phospho-ATR (T1989) (D5K8W) rabbit mAb (1:250, Cell Signaling Technology #30632); phospho-CHK1 (S317) rabbit mAb (1:250, Abcam #ab278717); FANCD2 rabbit pAb (1:500, Novus Biologicals #100-182); Polκ mouse mAb (1:500, Abcam #ab57070; mouse mAb is discontinued and #ab115625 goat pAb is recommended by manufacturer as a suitable replacement); β-tubulin (D3U1W) mouse mAb (1:1000, Cell Signaling Technology #86298); Ki67 sheep pAb (1:200, R&D Systems #AF7617). Secondary antibodies in this study include goat anti-rabbit, goat anti-mouse or donkey anti-sheep IgG secondaries linked to Alexa Fluor 488 (1:500, Thermo Fisher #A-11034, #A-11001), Alexa Fluor 546 (1:500, Thermo Fisher #A-11010, #A-11003, #A-21098), Alexa Fluor 647 (1:500, Thermo Fisher #A-21245, #A-21235), or HRP (1:3000, Cell Signaling Technology #7074, #7076). Unless otherwise noted, all primary and secondary antibody solutions were prepared in 3% BSA blocking buffer at the indicated dilutions.

### Western blotting

Cell suspensions were pelleted and lysed in RIPA lysis buffer supplemented with reducing agent and 1x phosphatase and protease inhibitor (MilliporeSigma #4906845001, #5892970001). Lysates were sheared with a 1cc U-100 insulin syringe (Becton Dickinson, #329424) and heated at 95 °C for 10 min. Proteins were separated by Bolt 4-12% Bis-Tris Plus gel (Thermo Fisher, NW04125BOX) and transferred to a PVDF membrane (Merck Millipore, #IPFL00010). The membrane was incubated in 3% BSA (GoldBio, #A-421-250) supplemented with 0.1% Tween-20 (Thermo Fisher, #9005-64-5) at room temperature for 2 h before overnight incubation with antibodies against γ-H2AX (1:400), FANCD2 (1:500) and β-tubulin (1:1000). The membrane was then washed for 5 min with PBS supplemented with 0.1% Tween-20 three times and then incubated with anti-rabbit or anti-mouse IgG, HRP-linked secondary antibodies for 1 h. The chemiluminescent signals were detected on an Azure C600 from Azure Biosystems and images were analyzed in Fiji.

### Immunofluorescence imaging and processing

Cells were seeded on a glass-bottom 96-well plate coated with collagen (1:50, Advanced Biomatrix #5005) 24 h prior to drug treatment. Seeding densities for 72 h and 96 h fixed-cell experiments were optimized to maintain cell health at time of seeding and to prevent contact inhibition by multi-day time points: A375, 1,000 cells/well; HT29, 2,000 cells/well; PC9, 2,000 cells/well; SW837, 7,500 cells/well; H358, 3,000 cells/well. Following treatments, cells were fixed with 4% paraformaldehyde for 10 min. For immunofluorescence, a standard protocol was used: cells were blocked in 3% BSA solution for 1 h at room temperature, permeabilized with 0.1% Triton X-100 at 4 °C for 15 min, incubated with primary antibodies overnight at 4 °C, and incubated with secondary antibodies for 2 h at room temperature. Where applicable, cells were stained with Hoechst 33342 (1:10,000) for 10 min to visualize DNA content. Immunofluorescence imaging of most markers were performed on a Nikon Ti-E using a 10x/0.45 numerical aperture (NA) objective. Single-cell quantitation of the captured images was performed using custom MATLAB scripts, where a nuclear mask was generated using the Hoechst 33342 signal, and each segmented nuclei was then probed for other signal intensities. Immunofluorescence imaging of γ-H2AX and FANCD2 stained cell cultures were performed on a PerkinElmer Opera Phenix high-content screening system with a 40x/1.0 NA water objective, acquired and analyzed via Harmony software and custom MATLAB scripts.

### Formalin-fixed paraffin-embedded (FFPE) tissue staining

Tissue sections from BRAF^V600E^ melanoma patients MB2282 and MB3022 were obtained from the Center for Rare Melanomas at Colorado Anschutz Medical Campus. Tissues were biopsied by the Center, appropriately deidentified to allow the samples to be used for research purposes, processed into FFPE blocks, cut, mounted, and sent to our lab for immunostaining. FFPE sections were then processed for immunofluorescence staining, as described by a standard protocol (*61*). Briefly, FFPE sections were washed in 3 rounds of SafeClear (xylene alternative, Fisher Scientific # 23-314629), followed by 3 rounds of ethanol. Heat-induced epitope retrieval was carried out by boiling slides in sodium citrate solution for 10 min (10 mM sodium citrate, 0.05% Tween 20, pH 6.0). Tissues were then blocked in 1x Carbo-Free blocking solution (Vector Laboratories #SP-5040-125) with 0.1% Triton X-100 for 2 h at room temperature, incubated with primary antibodies overnight at 4 °C, and incubated with secondary antibodies (1:500) and Hoechst 33342 (1:5,000) for 2 h at room temperature. Before and after antibody incubations, tissues were washed three times with PBS-T on a shaker for 5 min. Immunofluorescence imaging of tissue sections were performed on a Nikon Ti-E using a 20x/0.75 NA objective. Cells were scored for Ki67-positive status and γ-H2AX mean nuclear intensity was measured using custom MATLAB scripts.

### EdU incorporation

To identify cells actively synthesizing DNA, cells were pulsed with 10 μM of the synthetic nucleotide EdU (alkyne-conjugated) at 37 °C for 15 minutes prior to fixation with 4% paraformaldehyde. The EdU was visualized by click chemistry with azide-linked fluorescent dyes as described in the manufacturer’s protocol (Thermo Fisher, #C10340 and #C10641). Cells were then twice washed with PBS and blocked with 3% BSA for 1 h at room temperature to prepare for further immunostaining.

### Flow cytometry apoptosis assay

Cells were seeded in 12-well plates (Corning, #3513), at 10^5^ cells/well, 24 h prior to drug treatments. Wells were treated in triplicate with various doses and combinations of MAPK pathway inhibitors used in the study for 4 or 7 days. Etoposide (10 μM) was used as a positive control for apoptosis. After treatment, non-adherent cells were first harvested by pipetting, adherent cells were harvested by trypsinization, and these two populations were then combined. Cell suspensions were centrifuged and resuspended in calcium-rich binding buffer provided by the apoptosis staining kit (Abcam, #ab14085) to reach ~10^6^ cells/mL. Live suspensions were stained with both Annexin V-FITC (1:100) and propidium iodide, PI (1:100). Single-cell fluorescent signals were acquired on a BD FACSCelesta flow cytometer equipped with 488 and 561 nm lasers. By convention, cells were gated in FlowJo to remove debris and doublets. Annexin V-FITC and PI values were plotted as a bivariate scatter and etoposide-determined quadrant gating was applied to all plots to reach final apoptotic population percentages (see fig. S2).

### Live-cell time-lapse imaging and single-cell tracking

Cells were seeded on a glass-bottom 96-well plate coated with collagen 24 h prior to the start of imaging. Movie images were taken on a Nikon Ti-E using a 10x/0.45 NA objective (for imaging PC9, HT29, and A375 cells expressing DHB-mTurquoise2) or 20x/0.75 NA objective (for SW837 DHB-mTurquoise2 imaging, or for A375 FANCD2-mCitrine imaging) with appropriate filter sets at a frequency of 15 min per frame (for multi-day movies) or 4 min per frame (for shorter movies of mitotic intervals). Cells were maintained in a humidified incubation chamber at 37 °C with 5% CO2. Cells were imaged in phenol red-free full-growth media. For times of drug addition or siRNA treatments, the movie was paused to allow for treatment-containing media, and then imaging continued. For the treatments spanning multiple days, drug was refreshed 48 h after the initial drug addition time point. The drug refreshment was performed by exchanging half of the total media in each well to avoid cell loss during pipetting.

Multi-day movies were tracked using established pipelines (*62, 63*), as previously described (*24*). The open access tracking codes are available for download from GitHub in the referenced literature. Cell tracks were manually verified such that only cells correctly tracked during the entire movie were kept for downstream analysis. CDK2 activity was read out as the cytoplasmic:nuclear (C/N) ratio of the DHB signal, as previously described (*32*), where nuclear mean was used and the cytoplasmic component was calculated as the mean of the top 50th percentile of a ring of pixels outside of the nuclear mask. Cell-cycle entry points are defined as points where CDK2 activity begins to rise. The point of anaphase is defined when the H2B-based nuclear mask is divided to give rise to two new distinctly segmented bodies. Escapees are defined as cells with at least 10 h of drug-induced CDK2-low quiescence before cell-cycle re-entry. Non-escapees are defined as cells that never emerge from drug-induced CDK2-low quiescence by the end of the imaging period. For CDK2 activity heatmaps, escapees and non-escapees were plotted separately and cells within each category were sorted by the time of tracks ending due to cell death (characterized by nuclear shrinkage, dissolution, or explosion). The plots were then recombined. A375 cells expressing H2B-mIFP, DHB-mCherry, and FANCD2-mCitrine were tracked for CDK2 activity and verified, and FANCD2-mCitrine nuclear foci were quantified by automating each cell’s local maxima per frame in Fiji. Short-interval movies of cells that had been pre-treated with indicated drugs for indicated times were manually scored for mitotic durations, defined as the time of nuclear envelope breakdown (NEBD, defined by loss of DHB compartmentalization), accompanied shortly thereafter by chromatin condensation, to the time of anaphase. Since these movies contained 4 min frame intervals, the mitotic duration is approximated at this time resolution.

### Dose-response quantification

Dose-response curve fits for each cell line’s cell count after treatments were calculated using GraphPad Prism (v8.3). All cell line dose-response curves were fit using the following standard inhibitory Hill function.

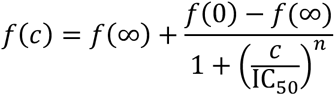

Here, *f*(*c*) is the fraction of cells in the population at drug concentration *c*; *f*(0) and *f*(∞) are the fractions at no drug and at the maximal tested drug concentration, respectively; IC_50_ is the half-maximal inhibitory concentration of the drug; and *n* is the Hill coefficient. *f*(*c*), *f*(0), and *f*(∞) were obtained from experiments; and the values of IC_50_ and *n* were obtained from fitting. The biphasic shape of the A375 dose-response curves was fitted by the summation of two sigmoidal inhibitory curves (employed in GraphPad Prism as a biphasic fit function).

### Long-term cell growth assay

For the experiments in Fig. 5A and fig. S8A, PC9, SW837 and A375 cells expressing H2B-mIFP nuclear marker were seeded into a 96-well plate 24 h prior to the start of treatments, and the same wells were tracked for the duration of the experiment. Drug media was refreshed every 4 days throughout the 57-day duration. Entire well images were captured on a PerkinElmer Opera Phenix (5X objective). Analysis was performed by automated approximation of H2B-mIFP nuclei at each time point and normalizing to cell counts at day 0.

### Patient survival analyses

Kaplan-Meier survival analyses are based on data generated from the Harmonized Cancer Datasets within the NCI Genomic Data Commons (GDC) research database (as of April 28, 2022; https://portal.gdc.cancer.gov) (*64*). The cases included in survival analyses are broken down by grouping shown (fig. S9A), where patients with tumours of all tissue origins were selected based on the condition that they harbor ≥ 1 EGFR, KRAS, or BRAF mutation. These cases were subcategorized based on the tumour sample’s wild-type TP53 status or by harboring ≥ 1 TP53 mutation. Cohort comparisons between determined subgroups containing wild-type or mutant forms of the indicated genes (see fig. S9A for FA gene list) were generated in the GDC portal and directly exported. All *p*-values regarding survival plots were determined by log-rank test.

### Statistics

For violin plots used throughout the paper, thick lines represent the median values unless otherwise indicated, and thin bars above and below each median represent the interquartile ranges of the distribution. The full distributions are displayed by the full range of the violin shape, with the width along the violin corresponding with the value frequency. Statistical tests were performed using GraphPad Prism. Significance levels are reported as *p* values ≤ 0.05 (*), 0.01 (**), 0.001 (***) and 0.0001 (****) with corresponding star notations. Throughout, ‘ns’ denotes no statistical significance (*p* > 0.05).

## Supplementary Materials

**Fig. S1.** EGFR-, KRAS-, and BRAF-mutant cancer cell types can survive and cycle through supraphysiological doses of MAPK pathway inhibitors.

**Fig. S2.** Multiple cancer cell types treated with multiple targeted MAPK pathway inhibitors experience incomplete apoptosis.

**Fig. S3.** Cancer cells slowly cycling through MAPK pathway inhibition display changes in cell-cycle phase lengths.

**Fig. S4.** Cancer cells slowly cycling through MAPK pathway inhibition display longer times spent in prophase, metaphase, anaphase, and telophase, collectively.

**Fig. S5.** Cells escaping MAPK inhibition mount DNA replication stress responses that allow for cytoprotective effects.

**Fig. S6.** Cancer cells escaping MAPK pathway inhibition rely on increased FANCD2 nuclear recruitment.

**Fig. S7.** MAPK-mutant cancer cells display persistent nuclear Polκ translocation after MAPK pathway inhibitor treatments.

**Fig. S8.** Cells cycling amid extended treatment to MAPK pathway inhibition rely on an intact DNA damage response.

**Fig. S9.** Clinical cases of *EGFR*-, *KRAS*-, or *BRAF*-mutant cancers display poorer prognoses when FA pathway is intact, especially in a *TP53*-mutant background.

**Video S1.** Example of an SW837 escapee cell tracked for 4 days after 100 nM AMG510 addition.

**Video S2.** PC9, H358, and A375 cells completing mitosis under MAPK inhibition. CDK2 sensor expression shown for each cell line and condition.

**Video S3.** SW387 cells completing mitosis under KRAS inhibition. CDK2 sensor shown.

**Video S4.** FANCD2 foci appearance in an A375 cell (clone 7) escaping dabrafenib treatment. Time stamp is relative to drug addition.

**Video S5.** Dabrafenib-treated A375 cells challenged with FANCD2 depletion. CDK2 sensor expression shown for each condition.

## Acknowledgements and funding

We thank members of the Spencer lab for general help and discussion. Dr. Matthew Hellman, Dr. Helena Yu, and Dr. Daniel Durocher for conversations and comments on our data. Dr. Kasey Couts and Robb Van Gulick at the Center for Rare Melanomas at Colorado Anschutz Medical Campus for providing patient tissue sections. Dr. Jean-Yves Masson at Laval University Cancer Research Center for the pLenti-FANCD2-mCitrine plasmid. Theresa Nahreini for her assistance in cell sorting in the Cell Culture and Flow Cytometry core facilities. The BD FACSCelesta cytometer and BD FACSAria Fusion cell sorter are supported by NIH Grant S10OD021601. Some imaging work was performed at the BioFrontiers Institute Advanced Light Microscopy Core (RRID: SCR_018302) with the help of Joseph Dragavon. The PerkinElmer Opera Phenix is supported by NIH grant 1S10OD025072. The results of Kaplan-Meier plots are based on data generated from the Harmonized Cancer Datasets within the Genomic Data Commons (GDC) research database (accessed April 28, 2022): https://portal.gdc.cancer.gov. This work is supported by a NIH NRSA training grant awarded to T.E.H. (T32CA174648), a Damon Runyon-Rachleff Innovation Award (RUNYON-68-21) to S.L.S., and a Mark Foundation Emerging Leader Award (MFCR-AWD-20-08-0164) to S.L.S.

## Author contributions

T.E.H. conducted most experiments, analyses, data interpretation and manuscript preparation. C.Y. helped develop the FANCD2-mCitrine cell lines and assisted with experiments. V.N. and C.R.I. assisted with western blot and fixed-cell immunofluorescence experiments. S.L.S. conceived the project, suggested the experiments, interpreted the data, and wrote the manuscript with T.E.H.

## Supplementary Materials

**Fig. S1.**
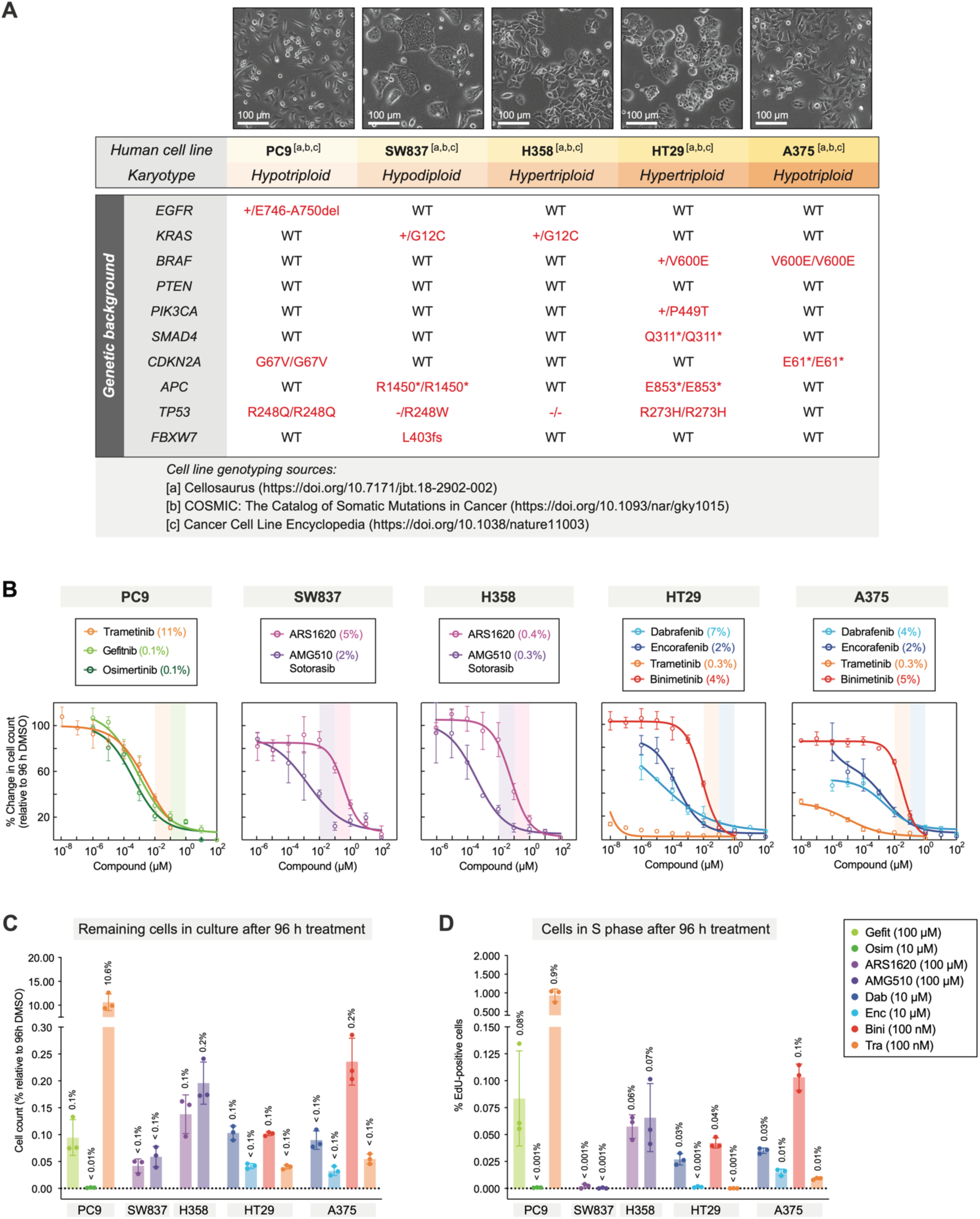
EGFR-, KRAS-, and BRAF-mutant cancer cell types can survive and cycle through supraphysiological doses of MAPK pathway inhibitors. **(A)** Genotype of each human cancer cell line identified as reported by indicated databases. **(B)** Dose-response profiling of MAPK-mutant cancer cell lines to their corresponding effective inhibitors. Inhibitory growth response is normalized relative to 96 h DMSO condition. Shaded bars cover dose ranges commonly used in cell culture models. Reported % value in plot legends corresponds to the measured value remaining at the highest concentration tested. Mean ± std of 3 replicate wells. **(C and D)**, Profiling of cancer cell responses to supraphysiological doses of MAPK inhibitors. Normalized cell count (C) and % EdU+ cells (D) were collected by fluorescence microscopy and custom MATLAB image processing. Mean ± std of 3 replicates.

**Fig. S2.**
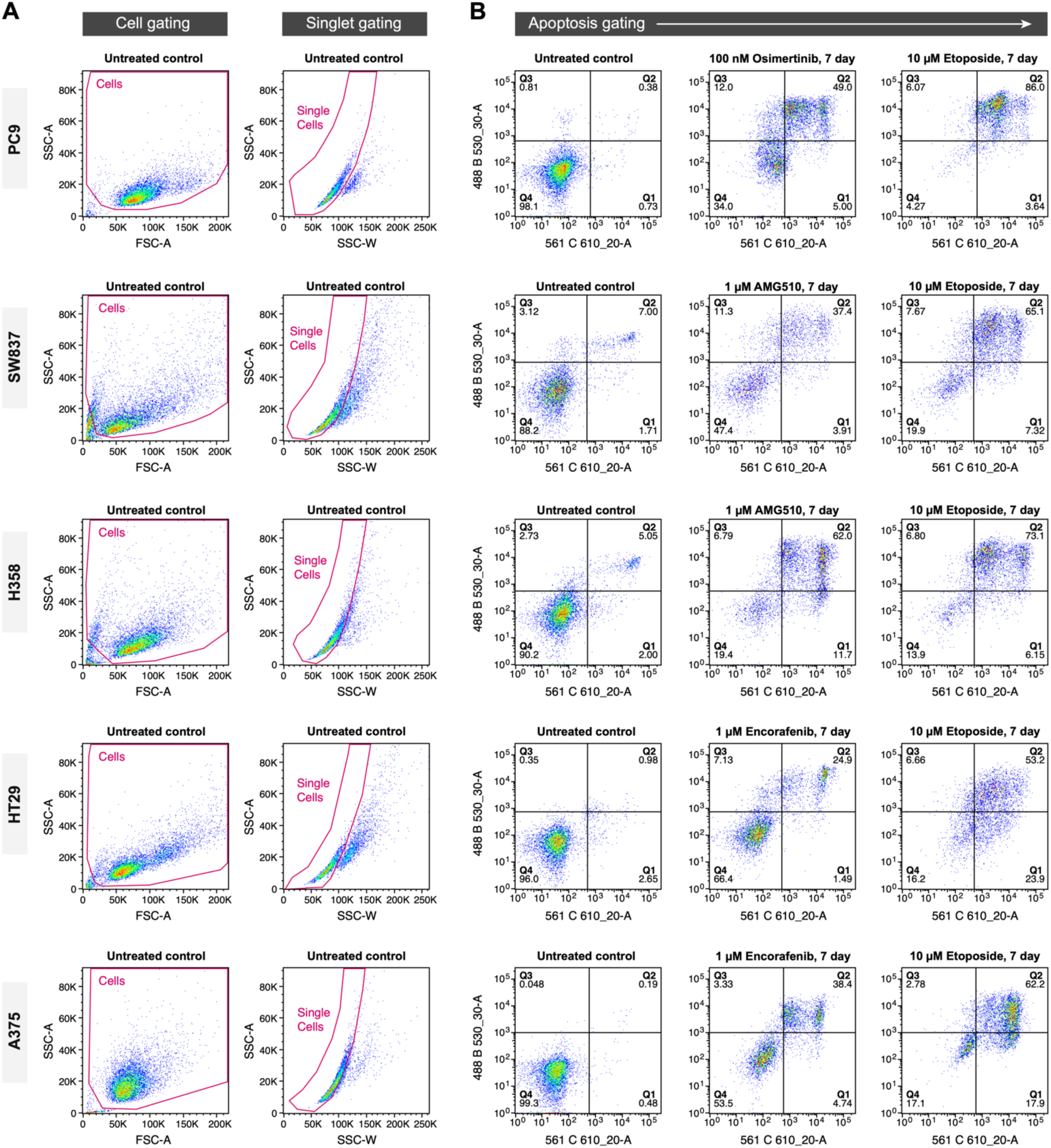
Multiple cancer cell types treated with multiple targeted MAPK pathway inhibitors experience incomplete apoptosis. **(A)** Flow cytometry collection of indicated call lines (n = 10,000 ungated events per replicate) and conventional single-cell gating scheme. FSC-A *vs.* SSC-A gating determines cellular events (small debris events excluded), and SSC-W *vs.* SSC-A gating determines singlet events (doublets excluded). **(B)** Apoptotic gating of single-cell events for all cell lines after indicated treatments, as determined by Annexin V staining (488 B 530_30 A) and propidium iodide (561 C 610_20-A). Etoposide is used as a positive control for a well-known extreme apoptotic response. Percent of cells in Q2 quadrant is reported in Fig. 1D.

**Fig. S3.**
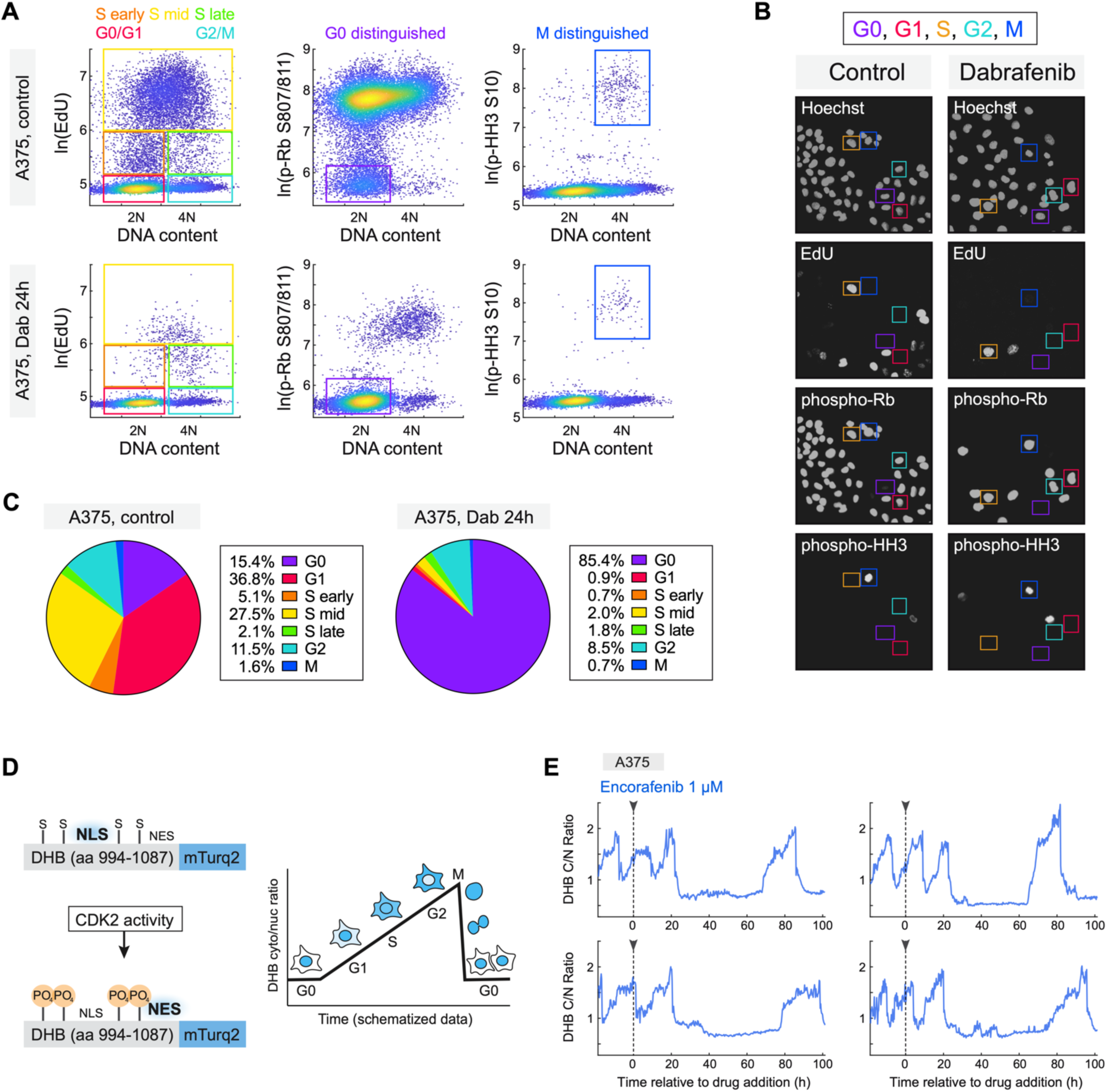
Cancer cells slowly cycling through MAPK pathway inhibition display changes in cell-cycle phase lengths. **(A)** Gated density scatters of fluorescence intensity measurements in A375 cells treated for 0 h or 24 h with 1 µM dabrafenib. *Left*, Hoechst intensity (DNA content) plotted against EdU incorporation (DNA synthesis) to determine cells in indicated cell cycle phases (G0/G1, S early, S mid, S late, and G2/M). *Middle*, DNA content vs. phospho-Rb (S807/811) to mark cell cycle commitment; G0/G1-gated cells are further gated into proper G0 and G1 designated populations as indicated. *Right*, DNA content vs. phospho-HH3 (S10) to mark mitotic entry; G2/M-gated cells are further gated into proper G2 and M designated populations as indicated. **(B)** Representative images of the imaging experiment quantified in (A). **(C)** Cell-cycle phase summary of A375 cells after 24 h of indicated treatments. **(D)** Schematic of DHB-based CDK2 sensor, adapted from Spencer et al. (2013). **(E)** Representative single-cell DHB C/N traces of A375 cells escaping from BRAF inhibition.

**Fig. S4.**
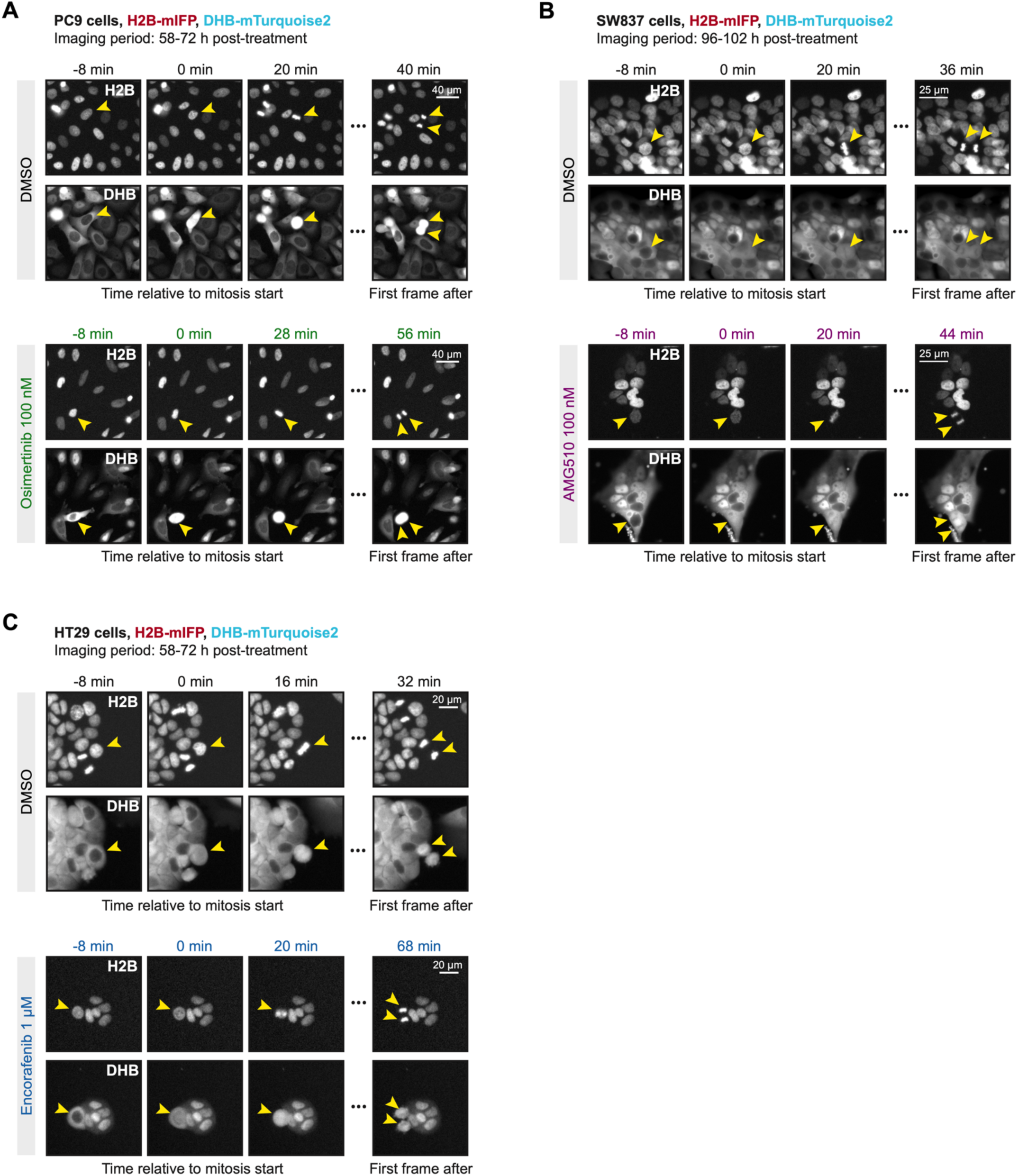
Cancer cells slowly cycling through MAPK pathway inhibition display longer times spent in prophase, metaphase, anaphase, and telophase, collectively. **(A to C)** Film strip of H2B and CDK2 activity sensor in PC9, SW837 or HT29 cells completing mitosis under DMSO treatment or MAPK inhibition.

**Fig. S5.**
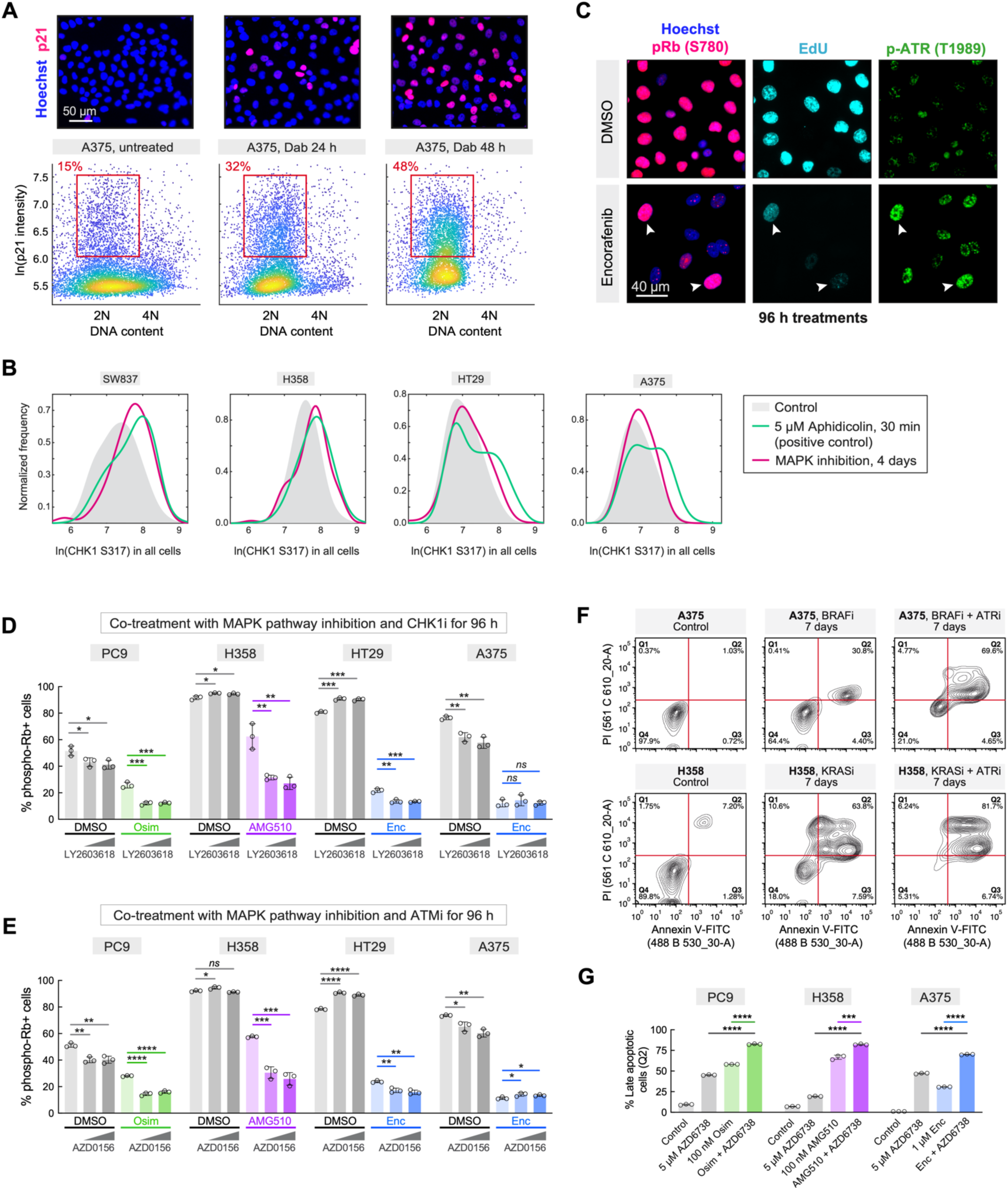
Cells escaping MAPK inhibition mount DNA replication stress responses that allow for cytoprotective effects. **(A)** Representative images (*top*) and density scatter plots (*bottom*) of DNA content and p21 immunofluorescence intensity in A375 cells treated with BRAF inhibition for 0, 24, and 48 h. The red gate represents 2N cells with high p21; percentage of population within the gate is displayed. **(B)** Histograms of phospho-CHK1 (S317) in SW837, H358, HT29, and A375 cells after indicated treatments. Cells shown here are ungated, whereas the histograms shown in main Fig. 3E have been gated on EdU-positive cells. Aphidicolin is used here as a positive control to confirm p-CHK1 (S317) staining. **(C)** Representative immunofluorescence images of phospho-ATR (T1989) in A375 cells escaping BRAF inhibition (cycling status displayed by phospho-Rb and EdU intensity). **(D and E)** Quantification of cycling cells (determined by phospho-Rb S807/811 immunofluorescence) after 96 h of indicated treatment combinations with CHK1 inhibition or ATM inhibition. CHK1i (LY603618) doses: 1 nM, 10 nM. ATMi (AZD0156) doses: 10 nM, 100 nM. Mean ± std of 3 replicates; *p*-value determined by unpaired t-test. **(F and G)** Flow cytometric apoptosis assay in MAPK-mutant cell lines after the indicated 7-day treatments. (F) Representative contour plots of A375 and H358 cells progressing to the final apoptotic fate (quadrant Q2) after MAPK inhibition and ATR inhibition. (G) Quantification of Q2 across indicated cell lines and treatments. Mean ± std of 3 replicates; *p*-value determined by unpaired t-test.

**Fig. S6.**
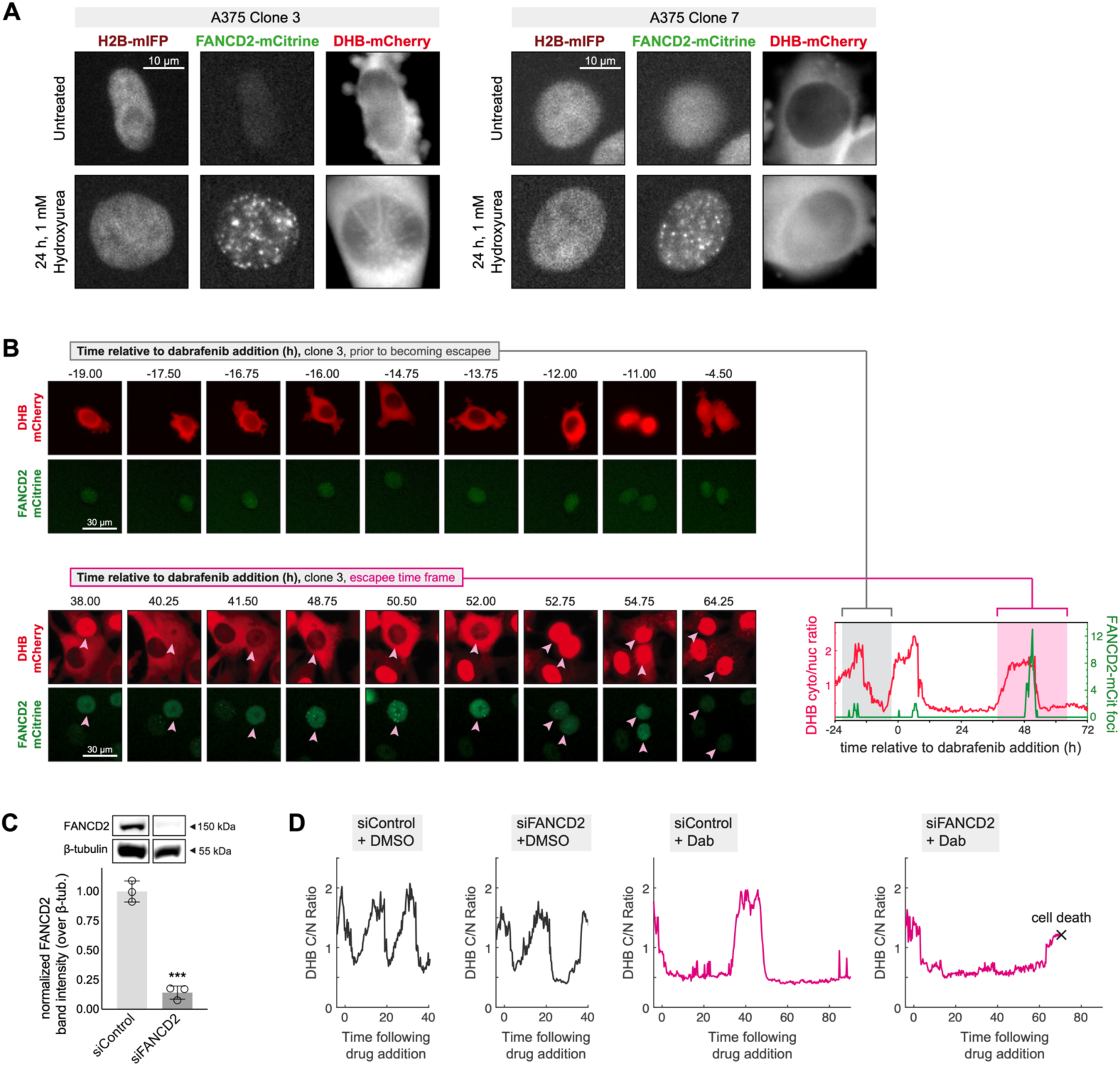
Cancer cells escaping MAPK pathway inhibition rely on increased FANCD2 nuclear recruitment. **(A)** Validation of FANCD2-mCitrine expressing clonal cell lines using hydroxyurea treatment as a positive control. Representative cells in S/G2 phases of the cell cycle (marked by cytoplasmic DHB-mCherry intensity) display many FANCD2-mCitrine nuclear foci after 24 h treatments, a known effect of hydroxyurea. **(B)** Film strips of FANCD2-mCitrine and DHB-mCherry in A375 cells under indicated times of dabrafenib treatment. Arrowheads follow a cell progressing through an entire cell cycle and experiencing FANCD2-mCitrine nuclear foci while under dabrafenib treatment. Corresponding DHB C/N ratio and FANCD2 foci traces for the cell lineage prior to becoming an escapee and afterward are matched to the film strip time frames, shown in the righthand plot. **(C)** Validation of siRNA knockdown of FANCD2 by western blotting after 24 h; β-tubulin is used as a loading control. FANCD2 depleted by ~90%. Mean ± std of 3 replicates; *p*-value determined by unpaired t-test. **(D)** Single-cell traces of CDK2 activity over time in cells treated with combination treatments. The transfection mix was added to the cells 4 h prior to the time of drug treatment and removed 4 h after the time of drug treatment.

**Fig. S7.**
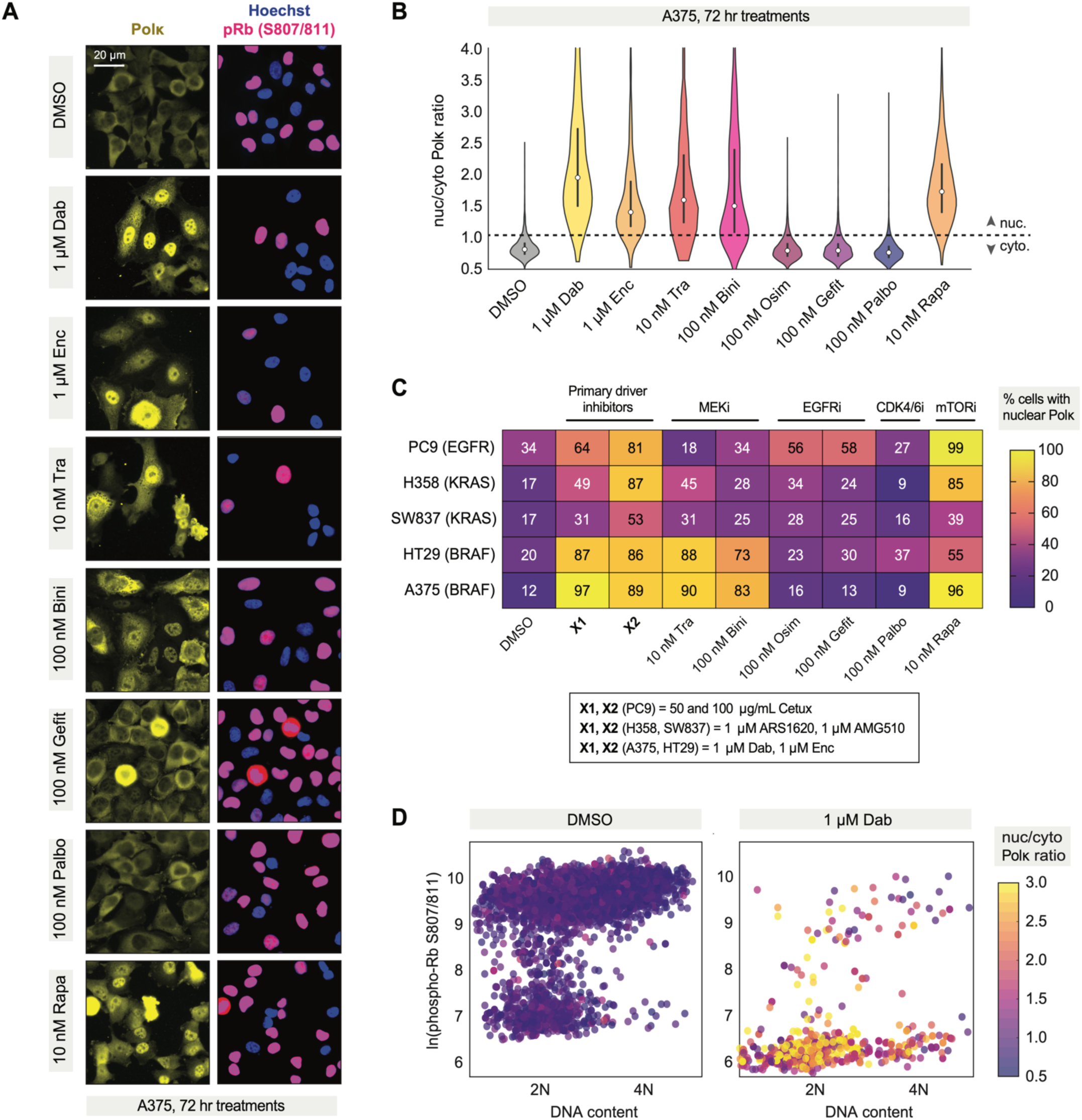
MAPK-mutant cancer cells display persistent nuclear Polκ translocation after MAPK pathway inhibitor treatments. **(A)** Representative immunofluorescence images of cells after 72 h of indicated treatments, stained for DNA content, phospho-Rb, and polymerase kappa (Polκ). The mTOR inhibitor rapamycin is used as a positive control for this translocation effect. **(B)** Quantification of nuc/cyto Polκ intensity fluorescence intensity after 72 h indicated treatments, displayed as violin distributions. Dashed line marks the 1:1 nuc/cyto ratio line. **(C)** Heat map of % cells with nuclear Polκ after 72 h of indicated MAPK pathway inhibitors, palbociclib, or rapamycin (positive control for nuclear translocation). Data are presented for the 5 different MAPK-mutant cell lines used throughout this study. **(D)** Scatter plots of DNA content and p-Rb in A375 cells treated with 72 h DMSO or BRAF inhibition. Each data point is color-coded to represent nuc/cyto Polκ intensity, as indicated by the colormap.

**Fig. S8.**
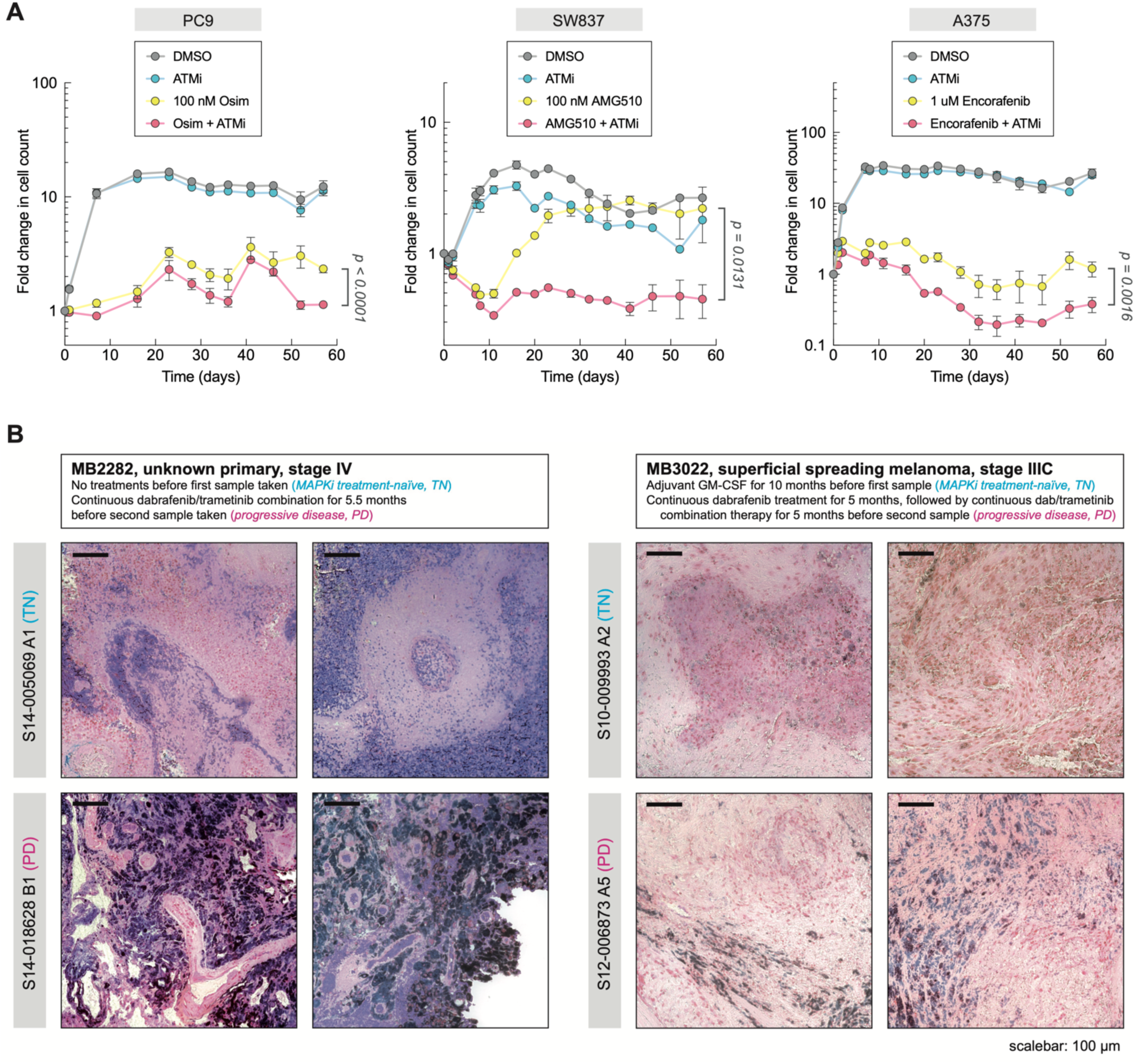
Cells cycling amid extended treatment to MAPK pathway inhibition rely on an intact DNA damage response. **(A)** Long-term culture of MAPK-mutant cell lines with indicated MAPK inhibitors and clinical ATM inhibitor (500 nM AZD0156). Cell count was quantified longitudinally by imaging H2B-mIFP fluorescence. Mean ± std of 4 replicate wells; *p*-value determined by unpaired t-test of final time point. **(B)** MB2282 and MB3022 patient treatment information prior to reaching a state of progressive disease. Representative haematoxylin and eosin staining of treatment-naïve (TN) and progressive disease (PD) tissue samples longitudinally biopsied from each patient.

**Fig. S9.**
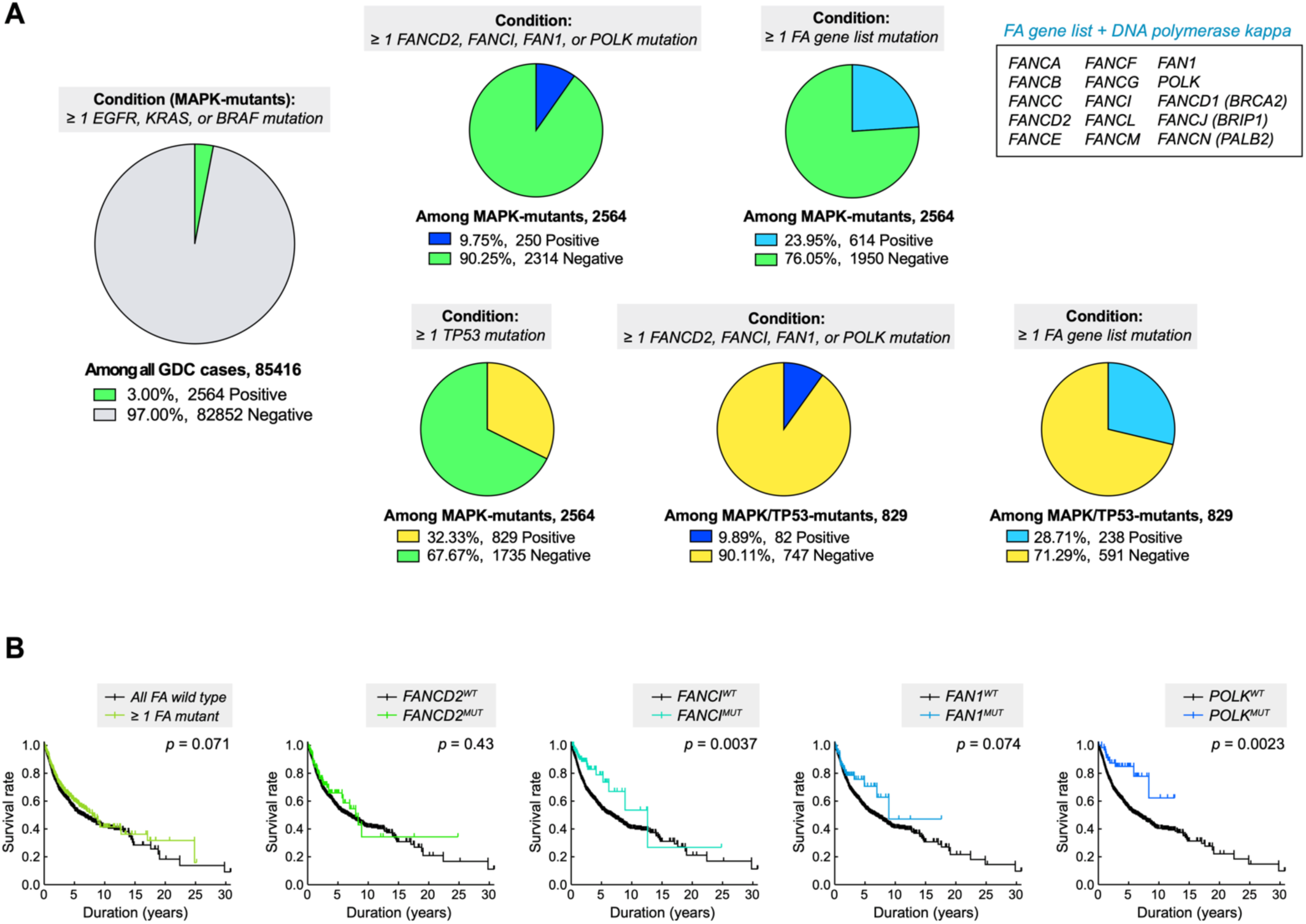
Clinical cases of *EGFR*-, *KRAS*-, or *BRAF*-mutant cancers display poorer prognoses when FA pathway is intact, especially in a *TP53*-mutant background. **(A)** Patient population data generated from the Harmonized Cancer Datasets within the Genomic Data Commons (GDC) research database (as of April 28, 2022; https://portal.gdc.cancer.gov). Patients were included with tumours harboring ≥ 1 EGFR, KRAS, or BRAF mutation initially, and subcategorized thereafter using the indicated conditions. Populations represent tumours of all tissue origins. Variants observed for FA gene list were 82% missense mutations, 12% stop-gain mutations, 6% frameshift mutations. While stop-gain mutations often result in loss-of-function variants, additional study is required to determine specific changes in protein function relative to each variant observed in the GDC database. **(B)** Kaplan-Meier survival analyses based on data generated from the GDC research database. Patients were included with tumours harboring ≥ 1 EGFR, KRAS, or BRAF mutation. In contrast to main Fig. 5D, plots shown here possess wild-type TP53. Cohort comparisons were made between tumour subgroups containing wild-type or mutant forms of the indicated genes. Left-most plot examines a cohort that has ≥ 1 mutation among the list of FA-associated genes from (A). Displayed *p*-values were determined by log-rank test.

## REFERENCES

1. Guo YJ, Pan WW, Liu SB, Shen ZF, Xu Y, Hu LL (2020) ERK/MAPK signalling pathway and tumorigenesis. Experimental and Therapeutic Medicine, 19(3), 1997–2007. https://doi.org/10.3892/etm.2020.8454

2. Zhang YL, Yuan JQ, Wang KF, Fu XH, Han XR, Threapleton D, Yang ZY, Mao C, Tang JL (2016) The prevalence of EGFR mutation in patients with non-small cell lung cancer: a systematic review and meta-analysis. Oncotarget, 7(48), 78985–78993. https://doi.org/10.18632/oncotarget.12587

3. Prior IA, Hood FE, Hartley JL (2020) The Frequency of Ras Mutations in Cancer. Cancer Research, 80(14), 2969–2974. https://doi.org/10.1158/0008-5472.CAN-19-3682

4. Seppälä TT, Böhm JP, Friman M, Lahtinen L, Väyrynen VM, Liipo TK, Ristimäki AP, Kairaluoma MV, Kellokumpu IH, Kuopio TH, Mecklin JP (2015) Combination of microsatellite instability and BRAF mutation status for subtyping colorectal cancer. British Journal of Cancer, 112(12), 1966–1975. https://doi.org/10.1038/bjc.2015.160

5. Davies H, Bignell GR, Cox C, Stephens P, Edkins S, Clegg S, Teague J, Woffendin H, Garnett MJ, Bottomley W, Davis N, Dicks E, Ewing R, Floyd Y, Gray K, Hall S, Hawes R, Hughes J, Kosmidou V, Menzies A, Mould C, Parker A, Stevens C, Watt S, Hooper S, Wilson R, Jayatilake H, Gusterson BA, Cooper C, Shipley J, Hargrave D, Pritchard-Jones K, Maitland N, Chenevix-Trench G, Riggins GJ, Bigner DD, Palmieri G, Cossu A, Flanagan A, Nicholson A, Ho JW, Leung SY, Yuen ST, Weber BL, Seigler HF, Darrow TL, Paterson H, Marais R, Marshall CJ, Wooster R, Stratton MR, Futreal PA (2002) Mutations of the BRAF gene in human cancer. Nature, 417(6892), 949–954. https://doi.org/10.1038/nature00766

6. Samatar AA, Poulikakos PI (2014) Targeting RAS-ERK signalling in cancer: promises and challenges. Nature Reviews. Drug Discovery, 13(12), 928–942. https://doi.org/10.1038/nrd4281

7. Cross DA, Ashton SE, Ghiorghiu S, Eberlein C, Nebhan CA, Spitzler PJ, Orme JP, Finlay MR, Ward RA, Mellor MJ, Hughes G, Rahi A, Jacobs VN, Red Brewer M, Ichihara E, Sun J, Jin H, Ballard P, Al-Kadhimi K, Rowlinson R, Klinowska T, Richmond GH, Cantarini M, Kim DW, Ranson MR, Pao W (2014) AZD9291, an irreversible EGFR TKI, overcomes T790M-mediated resistance to EGFR inhibitors in lung cancer. Cancer Discovery, 4(9), 1046–1061. https://doi.org/10.1158/2159-8290.CD-14-0337

8. Cheng Y, Tian H (2017) Current Development Status of MEK Inhibitors. Molecules, 22(10), 1551. https://doi.org/10.3390/molecules22101551

9. Rheault TR, Stellwagen JC, Adjabeng GM, Hornberger KR, Petrov KG, Waterson AG, Dickerson SH, Mook RA Jr, Laquerre SG, King AJ, Rossanese OW, Arnone MR, Smitheman KN, Kane-Carson LS, Han C, Moorthy GS, Moss KG, Uehling DE (2013) Discovery of Dabrafenib: A Selective Inhibitor of Raf Kinases with Antitumor Activity against B-Raf-Driven Tumors. ACS Medicinal Chemistry Letters, 4(3), 358–362. https://doi.org/10.1021/ml4000063

10. Hamid O, Cowey CL, Offner M, Faries M, Carvajal RD (2019) Efficacy, Safety, and Tolerability of Approved Combination BRAF and MEK Inhibitor Regimens for *BRAF*-Mutant Melanoma. Cancers, 11(11), 1642. https://doi.org/10.3390/cancers11111642

11. Canon J, Rex K, Saiki AY, Mohr C, Cooke K, Bagal D, Gaida K, Holt T, Knutson CG, Koppada N, Lanman BA, Werner J, Rapaport AS, San Miguel T, Ortiz R, Osgood T, Sun JR, Zhu X, McCarter JD, Volak LP, Houk BE, Fakih MG, O’Neil BH, Price TJ, Falchook GS, Desai J, Kuo J, Govindan R, Hong DS, Ouyang W, Henary H, Arvedson T, Cee VJ, Lipford JR (2019) The clinical KRAS(G12C) inhibitor AMG 510 drives anti-tumour immunity. Nature, 575(7781), 217–223. https://doi.org/10.1038/s41586-019-1694-1

12. Janes MR, Zhang J, Li LS, Hansen R, Peters U, Guo X, Chen Y, Babbar A, Firdaus SJ, Darjania L, Feng J, Chen JH, Li S, Li S, Long YO, Thach C, Liu Y, Zarieh A, Ely T, Kucharski JM, Kessler LV, Wu T, Yu K, Wang Y, Yao Y, Deng X, Zarrinkar PP, Brehmer D, Dhanak D, Lorenzi MV, Hu-Lowe D, Patricelli MP, Ren P, Liu Y (2018). Targeting KRAS Mutant Cancers with a Covalent G12C-Specific Inhibitor. Cell, 172(3), 578–589.e17. https://doi.org/10.1016/j.cell.2018.01.006

13. Yang A, Li M, Fang M (2021) The Research Progress of Direct KRAS G12C Mutation Inhibitors. Pathology Oncology Research, 27, 631095. https://doi.org/10.3389/pore.2021.631095

14. Chatterjee N, Bivona TG (2019) Polytherapy and Targeted Cancer Drug Resistance. Trends in Cancer, 5(3), 170–182. https://doi.org/10.1016/j.trecan.2019.02.003

15. Wu JY, Wu SG, Yang CH, Gow CH, Chang YL, Yu CJ, Shih JY, Yang PC (2008) Lung cancer with epidermal growth factor receptor exon 20 mutations is associated with poor gefitinib treatment response. Clinical Cancer Research, 14(15), 4877–4882. https://doi.org/10.1158/1078-0432.CCR-07-5123

16. Wagle N, Emery C, Berger MF, Davis MJ, Sawyer A, Pochanard P, Kehoe SM, Johannessen CM, Macconaill LE, Hahn WC, Meyerson M, Garraway LA (2011) Dissecting Therapeutic Resistance to RAF Inhibition in Melanoma by Tumor Genomic Profiling. Journal of Clinical Oncology. 29, 3085–3096. https://doi.org/10.1200/jco.2010.33.2312

17. Van Allen EM, et al. (2014) The Genetic Landscape of Clinical Resistance to RAF Inhibition in Metastatic Melanoma. Cancer Discovery. 4, 94–109. https://doi.org/10.1158/2159-8290.cd-13-0617

18. Shaffer SM, Dunagin MC, Torborg SR, Torre EA, Emert B, Krepler C, Beqiri M, Sproesser K, Brafford PA, Xiao M, Eggan E, Anastopoulos IN, Vargas-Garcia CA, Singh A, Nathanson KL, Herlyn M, Raj A (2017) Rare cell variability and drug-induced reprogramming as a mode of cancer drug resistance. Nature, 546(7658), 431–435. https://doi.org/10.1038/nature22794

19. Sharma SV, Lee DY, Li B, Quinlan MP, Takahashi F, Maheswaran S, McDermott U, Azizian N, Zou L, Fischbach MA, Wong KK, Brandstetter K, Wittner B, Ramaswamy S, Classon M, Settleman J (2010) A chromatin-mediated reversible drug-tolerant state in cancer cell subpopulations. Cell, 141(1), 69–80. https://doi.org/10.1016/j.cell.2010.02.027

20. Ramirez M, Rajaram S, Steininger RJ, Osipchuk D, Roth MA, Morinishi LS, Evans L, Ji W, Hsu CH, Thurley K, Wei S, Zhou A, Koduru PR, Posner BA, Wu LF, Altschuler SJ (2016) Diverse drug-resistance mechanisms can emerge from drug-tolerant cancer persister cells. Nature Communications. 7, 10690. https://doi.org/10.1038/ncomms10690

21. Hata AN, Niederst MJ, Archibald HL, Gomez-Caraballo M, Siddiqui FM, Mulvey HE, Maruvka YE, Ji F, Bhang HE, Krishnamurthy Radhakrishna V, Siravegna G, Hu H, Raoof S, Lockerman E, Kalsy A, Lee D, Keating CL, Ruddy DA, Damon LJ, Crystal AS, Costa C, Piotrowska Z, Bardelli A, Iafrate AJ, Sadreyev RI, Stegmeier F, Getz G, Sequist LV, Faber AC, Engelman JA (2016) Tumor cells can follow distinct evolutionary paths to become resistant to epidermal growth factor receptor inhibition. Nature Medicine. 22(3), 262–269. https://doi.org/10.1038/nm.4040

22. Fallahi-Sichani M, Becker V, Izar B, Baker GJ, Lin JR, Boswell SA, Shah P, Rotem A, Garraway LA, Sorger PK (2017) Adaptive resistance of melanoma cells to RAF inhibition via reversible induction of a slowly dividing de-differentiated state. Molecular Systems Biology. 13(1), 905. https://doi.org/10.15252/msb.20166796

23. Xue JY, Zhao Y, Aronowitz J, Mai TT, Vides A, Qeriqi B, Kim D, Li C, de Stanchina E, Mazutis L, Risso D, Lito P (2020) Rapid non-uniform adaptation to conformation-specific KRAS(G12C) inhibition. Nature. 577(7790), 421–425. https://doi.org/10.1038/s41586-019-1884-x

24. Yang C, Tian C, Hoffman TE, Jacobsen NK, Spencer SL (2021). Melanoma subpopulations that rapidly escape MAPK pathway inhibition incur DNA damage and rely on stress signalling. Nature Communications, 12(1), 1747. https://doi.org/10.1038/s41467-021-21549-x

25. Oren Y, Tsabar M, Cuoco MS, Amir-Zilberstein L, Cabanos HF, Hütter JC, Hu B, Thakore PI, Tabaka M, Fulco CP, Colgan W, Cuevas BM, Hurvitz SA, Slamon DJ, Deik A, Pierce KA, Clish C, Hata AN, Zaganjor E, Lahav G, Politi K, Brugge JS, Regev A (2021) Cycling cancer persister cells arise from lineages with distinct programs. Nature, 596(7873), 576–582. https://doi.org/10.1038/s41586-021-03796-6

26. Russo M, Crisafulli G, Sogari A, Reilly NM, Arena S, Lamba S, Bartolini A, Amodio V, Magrì A, Novara L, Sarotto I, Nagel ZD, Piett CG, Amatu A, Sartore-Bianchi A, Siena S, Bertotti A, Trusolino L, Corigliano M, Gherardi M, Lagomarsino MC, Di Nicolantonio F, Bardelli A (2019) Adaptive mutability of colorectal cancers in response to targeted therapies. Science. 366(6472), 1473–1480. https://doi.org/10.1126/science.aav4474

27. Russo M, Pompei S, Sogari A, Corigliano M, Crisafulli G, Puliafito A, Lamba S, Erriquez J, Bertotti A, Gherardi M, Di Nicolantonio F, Bardelli A, Cosentino Lagomarsino M (2022). A modified fluctuation-test framework characterizes the population dynamics and mutation rate of colorectal cancer persister cells. Nature Genetics, 54(7), 976–984. https://doi.org/10.1038/s41588-022-01105-z

28. Lukow DA, Sausville EL, Suri P, Chunduri NK, Wieland A, Leu J, Smith JC, Girish V, Kumar AA, Kendall J, Wang Z, Storchova Z, Sheltzer JM (2021) Chromosomal instability accelerates the evolution of resistance to anti-cancer therapies. Developmental Cell, 56(17), 2427–2439.e4. https://doi.org/10.1016/j.devcel.2021.07.009

29. Moser J, Miller I, Carter D, Spencer SL (2018) Control of the Restriction Point by Rb and p21. Proceedings of the National Academy of Sciences of the United States of America. 115(35), E8219–E8227. https://doi.org/10.1073/pnas.1722446115

30. Daigh LH, Liu C, Chung M, Cimprich KA, Meyer T (2018) Stochastic Endogenous Replication Stress Causes ATR-Triggered Fluctuations in CDK2 Activity that Dynamically Adjust Global DNA Synthesis Rates. Cell Systems, 7(1), 17–27.e3. https://doi.org/10.1016/j.cels.2018.05.011

31. Lemmens B, Hegarat N, Akopyan K, Sala-Gaston J, Bartek J, Hochegger H, Lindqvist A (2018) DNA Replication Determines Timing of Mitosis by Restricting CDK1 and PLK1 Activation. Molecular Cell, 71(1), 117–128.e3. https://doi.org/10.1016/j.molcel.2018.05.026

32. Spencer SL, Cappell SD, Tsai FC, Overton KW, Wang CL, Meyer T (2013) The proliferation-quiescence decision is controlled by a bifurcation in CDK2 activity at mitotic exit. Cell. 155(2), 369–383. https://doi.org/10.1016/j.cell.2013.08.062

33. Masamsetti VP, Low R, Mak KS, O’Connor A, Riffkin CD, Lamm N, Crabbe L, Karlseder J, Huang D, Hayashi MT, Cesare AJ (2019) Replication stress induces mitotic death through parallel pathways regulated by WAPL and telomere deprotection. Nature Communications, 10(1), 4224. https://doi.org/10.1038/s41467-019-12255-w

34. Minocherhomji S, Ying S, Bjerregaard VA, Bursomanno S, Aleliunaite A, Wu W, Mankouri HW, Shen H, Liu Y, Hickson ID (2015) Replication stress activates DNA repair synthesis in mitosis. Nature. 528(7581), 286–290. https://doi.org/10.1038/nature16139

35. Mah LJ, El-Osta A, Karagiannis TC (2010) gammaH2AX: a sensitive molecular marker of DNA damage and repair. Leukemia, 24(4), 679–686. https://doi.org/10.1038/leu.2010.6

36. Arora M, Moser J, Phadke H, Basha AA, Spencer SL (2017) Endogenous Replication Stress in Mother Cells Leads to Quiescence of Daughter Cells. Cell Reports. 19(7), 1351–1364. https://doi.org/10.1016/j.celrep.2017.04.055

37. Fong CS, Mazo G, Das T, Goodman J, Kim M, O’Rourke BP, Izquierdo D, Tsou MF (2016) 53BP1 and USP28 mediate p53-dependent cell cycle arrest in response to centrosome loss and prolonged mitosis. eLife, 5, e16270. https://doi.org/10.7554/eLife.16270

38. Saldivar JC, Cortez D, Cimprich KA (2017) The essential kinase ATR: ensuring faithful duplication of a challenging genome. Nature Reviews Molecular Cell Biology, 18(10), 622–636. https://doi.org/10.1038/nrm.2017.67

39. Lebrec V, Poteau M, Morretton JP, Gavet O (2022) Chk1 dynamics in G2 phase upon replication stress predict daughter cell outcome. Developmental Cell, 57(5), 638–653.e5. https://doi.org/10.1016/j.devcel.2022.02.013

40. Badra Fajardo N, Taraviras S, Lygerou Z (2022) Fanconi anemia proteins and genome fragility: unraveling replication defects for cancer therapy. Trends in Cancer, S2405-8033(22)00022-X. Advance online publication. https://doi.org/10.1016/j.trecan.2022.01.015

41. Lossaint G, Larroque M, Ribeyre C, Bec N, Larroque C, Décaillet C, Gari K, Constantinou A (2013) FANCD2 binds MCM proteins and controls replisome function upon activation of S phase checkpoint signaling. Molecular Cell. 51(5), 678–690. https://doi.org/10.1016/j.molcel.2013.07.023

42. Chen YH, Jones MJ, Yin Y, Crist SB, Colnaghi L, Sims RJ 3rd, Rothenberg E, Jallepalli PV, Huang TT (2015) ATR-mediated phosphorylation of FANCI regulates dormant origin firing in response to replication stress. Molecular Cell. 58(2), 323–338. https://doi.org/10.1016/j.molcel.2015.02.031

43. Kais Z, Rondinelli B, Holmes A, O’Leary C, Kozono D, D’Andrea AD, Ceccaldi R (2016) FANCD2 Maintains Fork Stability in BRCA1/2-Deficient Tumors and Promotes Alternative End-Joining DNA Repair. Cell Reports, 15(11), 2488–2499. https://doi.org/10.1016/j.celrep.2016.05.031

44. Madireddy A, Kosiyatrakul ST, Boisvert RA, Herrera-Moyano E, García-Rubio ML, Gerhardt J, Vuono EA, Owen N, Yan Z, Olson S, Aguilera A, Howlett NG, Schildkraut CL (2016) FANCD2 Facilitates Replication through Common Fragile Sites. Molecular Cell. 64(2), 388–404. https://doi.org/10.1016/j.molcel.2016.09.017

45. Okamoto Y, Iwasaki WM, Kugou K, Takahashi KK, Oda A, Sato K, Kobayashi W, Kawai H, Sakasai R, Takaori-Kondo A, Yamamoto T, Kanemaki MT, Taoka M, Isobe T, Kurumizaka H, Innan H, Ohta K, Ishiai M, Takata M (2018) Replication stress induces accumulation of FANCD2 at central region of large fragile genes. Nucleic Acids Research. 46(6), 2932–2944. https://doi.org/10.1093/nar/gky058

46. Niraj J, Caron MC, Drapeau K, Bérubé S, Guitton-Sert L, Coulombe Y, Couturier AM, Masson JY (2017) The identification of FANCD2 DNA binding domains reveals nuclear localization sequences. Nucleic Acids Research. 45(14), 8341–8357. https://doi.org/10.1093/nar/gkx543

47. Bhowmick R, Minocherhomji S, Hickson ID (2016) RAD52 Facilitates Mitotic DNA Synthesis Following Replication Stress. Molecular Cell. 64(6), 1117–1126. https://doi.org/10.1016/j.molcel.2016.10.037

48. Helbling-Leclerc A, Dessarps-Freichey F, Evrard C, Rosselli F (2019) Fanconi anemia proteins counteract the implementation of the oncogene-induced senescence program. Scientific Reports. 9(1), 17024. https://doi.org/10.1038/s41598-019-53502-w

49. Bourseguin J, Bonet C, Renaud E, Pandiani C, Boncompagni M, Giuliano S, Pawlikowska P, Karmous-Benailly H, Ballotti R, Rosselli F, Bertolotto C (2016) FANCD2 functions as a critical factor downstream of MiTF to maintain the proliferation and survival of melanoma cells. Scientific Reports. 6, 36539. https://doi.org/10.1038/srep36539

50. Tonzi P, Yin Y, Lee C, Rothenberg E, Huang TT (2018) Translesion polymerase kappa-dependent DNA synthesis underlies replication fork recovery. eLife, 7, e41426. https://doi.org/10.7554/eLife.41426

51. Temprine K, Campbell NR, Huang R, Langdon EM, Simon-Vermot T, Mehta K, Clapp A, Chipman M, White RM (2020) Regulation of the error-prone DNA polymerase Polκ by oncogenic signaling and its contribution to drug resistance. Science Signaling, 13(629), eaau1453. https://doi.org/10.1126/scisignal.aau1453

52. Miller I, Min M, Yang C, Tian C, Gookin S, Carter D, Spencer SL (2018) Ki67 is a Graded Rather than a Binary Marker of Proliferation versus Quiescence. Cell Reports. 24(5), 1105–1112.e5. https://doi.org/10.1016/j.celrep.2018.06.110

53. Zhu G, Pan C, Bei JX, Li B, Liang C, Xu Y, Fu X (2020) Mutant p53 in Cancer Progression and Targeted Therapies. Frontiers in Oncology, 10, 595187. https://doi.org/10.3389/fonc.2020.595187

54. Santana-Codina N, Roeth AA, Zhang Y, Yang A, Mashadova O, Asara JM, Wang X, Bronson RT, Lyssiotis CA, Ying H, Kimmelman AC (2018) Oncogenic KRAS supports pancreatic cancer through regulation of nucleotide synthesis. Nature Communications, 9(1), 4945. https://doi.org/10.1038/s41467-018-07472-8

55. Ali ES, Sahu U, Villa E, O’Hara BP, Gao P, Beaudet C, Wood AW, Asara JM, Ben-Sahra I (2020) ERK2 Phosphorylates PFAS to Mediate Posttranslational Control of De Novo Purine Synthesis. Molecular Cell, 78(6), 1178–1191.e6. https://doi.org/10.1016/j.molcel.2020.05.001

56. Petropoulos M, Champeris Tsaniras S, Taraviras S, Lygerou Z (2019) Replication Licensing Aberrations, Replication Stress, and Genomic Instability. Trends in Biochemical Sciences. 44(9), 752–764. https://doi.org/10.1016/j.tibs.2019.03.011

57. Bowry A, Kelly R, Petermann E (2021) Hypertranscription and replication stress in cancer. Trends in Cancer, 7(9), 863–877. https://doi.org/10.1016/j.trecan.2021.04.006

58. Roos WP, Thomas AD, Kaina B (2016) DNA damage and the balance between survival and death in cancer biology. Nature Reviews Cancer. 16(1), 20–33. https://doi.org/10.1038/nrc.2015.2

59. Baillie KE, Stirling PC (2021) Beyond Kinases: Targeting Replication Stress Proteins in Cancer Therapy. Trends in Cancer, 7(5), 430–446. https://doi.org/10.1016/j.trecan.2020.10.010

60. Ali M, Lu M, Ang HX, Soderquist RS, Eyler CE, Hutchinson HM, Glass C, Bassil CF, Lopez OM, Kerr DL, Falcon CJ, Yu HA, Hata AN, Blakely CM, McCoach CE, Bivona TG, Wood KC (2022) Small-molecule targeted therapies induce dependence on DNA double-strand break repair in residual tumor cells. Science Translational Medicine, 14(638), eabc7480. https://doi.org/10.1126/scitranslmed.abc7480

61. Zaqout S, Becker LL, Kaindl AM (2020) Immunofluorescence Staining of Paraffin Sections Step by Step. Frontiers in Neuroanatomy, 14, 582218. https://doi.org/10.3389/fnana.2020.582218

62. Cappell SD, Chung M, Jaimovich A, Spencer SL, Meyer T (2016) Irreversible APCCdh1 Inactivation Underlies the Point of No Return for Cell-Cycle Entry. Cell, 166, 167–180. https://doi.org/10.1016/j.cell.2016.05.077

63. Tian C, Yang C, Spencer SL (2020) EllipTrack: A Global-Local Cell-Tracking Pipeline for 2D Fluorescence Time-Lapse Microscopy. Cell Reports, 32(5), 107984. https://doi.org/10.1016/j.celrep.2020.107984

64. Grossman RL, Heath AP, Ferretti V, Varmus HE, Lowy DR, Kibbe WA, Staudt LM (2016) Toward a Shared Vision for Cancer Genomic Data. The New England Journal of Medicine, 375(12), 1109–1112. https://doi.org/10.1056/NEJMp1607591

